# Harnessing protein-folding algorithms to drug intrinsically disordered epitopes

**DOI:** 10.1101/2025.11.11.687846

**Authors:** Jakub Lála, Stefano Angioletti-Uberti

## Abstract

Due to their lack of a specific structure and dynamical nature, targeting of epitopes that are part of an intrinsically disordered region of a protein is a notoriously difficult task. Here, we describe a computational approach to overcome this problem, based on the use of a protein-folding algorithm and its confidence metrics within a Monte Carlo optimization pipeline to generate peptide-based binders. For different protein targets, we show by accurate free energy calculations that our approach is able to design peptides with binding free energies on the order of tens of *k*_*B*_*T*, i.e., with strengths comparable to covalent interactions. Direct observation of the bound complex through molecular simulations shows that the targeted epitope folds into structured domains with lowered thermal fluctuations upon binding, while remaining unstructured and dynamic in the unbound state, suggesting that the protein-folding algorithm must have learned the principles of induced (co-)folding. Given the ubiquitous presence of unstructured regions in proteins, our results suggest a potential pathway to design drugs targeting a large variety of previously untargetable epitopes, and open new possibilities for therapeutic intervention in diseases where disordered proteins play a key role.

**SIGNIFICANCE:** Small-molecule drugs that bind to a protein via a lock-and-key mechanism require the targeted epitope to form a well-structured, stable binding pocket, thereby preventing binding to unstructured regions. To overcome this limit, we present a general approach, based on a protein-folding algorithm, to find peptide sequences that induce the formation of such a binding interface when no pocket is normally present. In other words, we show how a protein-folding algorithm can be used to program epitope recognition by *induced folding* instead of rigid lock-and-key matching. In this way, we show that we can extend druggable epitopes to intrinsically disordered, dynamical regions of proteins.

## II. INTRODUCTION

Intrinsically disordered regions (IDRs) of proteins are regions that lack a stable secondary structure, e.g., an alpha-helix or a beta-sheet. These regions are highly dynamic and participate in transient but specific interactions with other biomolecules, making them essential for biological functions. In fact, despite (or, maybe, because of) this characteristic, IDRs play critical roles in cellular processes, including cell signaling, regulation, and molecular recognition.^1–3^ Recently, they have also been implicated in liquid-liquid phase transitions that are proving critical to understanding how cells respond to stressors.^4,5^ Compared to well-structured regions, which are amenable to many of the tools developed by structural biologists over the past decades, the dynamical nature of IDRs makes it difficult to characterize them, and thus to understand their behavior, complicating the targeting of these regions with small-molecule drugs. One of the central challenges is that the absence of a stable structure hinders the application of one of drug design’s most successful paradigms: the lock-and-key model. As a result, traditional drug design strategies such as docking,^6^ which rely on well-defined binding pockets and stable secondary structures, are intrinsically inadequate for IDRs. For this reason, epitopes within these regions have often been defined as undruggable.^7^ RNA is also a highly flexible molecule, and, for similar arguments, designing drugs to target this nucleic acid has proven a hard problem. It has been shown, however, that for targeting RNA, some drugs achieve strong and selective binding by stabilizing a specific configuration.^8,9^This mechanism suggests, at least in principle, a potential pathway toward designing binders for proteins’ IDRs. Considering the practically infinite design space available, finding a generic molecule that stabilizes a specific fold in a protein might seem a prohibitively large task. To simplify the problem, we restrict ourselves to protein-based binders. By doing so, we aim to exploit a known mechanism, induced (co-)folding, whereby one or both otherwise unstructured proteins undergo a coordinated transition into a structured conformation upon binding. This mechanism has long been invoked as a way to mediate protein-protein interactions.^10,11^ In fact, it has already been suggested that signaling through interacting IDRs exploits induced folding to achieve selective binding.^2^ Whether or not two unstructured chains of amino acids can co-fold upon finding each other will depend on the formation of structural patterns, e.g., hydrogen bonds or patches of opposite charge, that can stabilize such a folded structure, and lead to attractive interactions between the two molecules. We notice that the same principle equally applies when considering the interaction between two distant parts of the same protein chain. Based on this premise, we hypothesize that to understand how proteins fold into their native structures, protein-folding algorithms must have also learned about induced folding. If this hypothesis is correct, one should be able to use such algorithms to design binders by looking for protein sequences that induce folding of a target IDR.

### A. Binder design strategy and system description

For a binder consisting of *N* canonical amino acids, there are 20^*N*^ potential candidate sequences to consider, making a random search computationally intractable for all but the shortest sequences. Instead, here we use Monte Carlo optimization as a strategy for the directed exploration of the binder sequence space, driven by the minimization of an energy function that, crucially, contains as a key ingredient the predicted Local Distance Difference Test (pLDDT)^12,13^ of the targeted IDR, a quantity returned by modern protein-folding algorithms. This metric is central to our purpose because sequence-to-structure models, such as AlphaFold2^13^ and ESMFold,^14^ have been shown to indirectly predict the dynamical, disordered nature of a residue’s local environment through the pLDDT.^15,16^The reason for this correlation is readily explained. The pLDDT was originally designed to convey the algorithm’s confidence in the prediction of a local environment around a given residue. However, one can never truly decouple two qualitatively different cases: low pLDDT values can either be the result of a truly inaccurate prediction, or, alternatively, indicate that the algorithm cannot associate a specific structure to such a region, exactly because no dominant one is present. The latter behavior is exactly what is expected for a dynamic region, i.e., an IDR.

We exploit this observation to design protein sequences that can bind to a target IDR through induced folding in the following way (details in the Methods section), schematically represented in Figure 1. We start from a random amino acid sequence for our binder and make single point mutations. We refold the protein in the presence of the mutated binder, and evaluate an energy that depends on the structure associated with the new sequence. This energy contains a term that increases if the proposed mutation decreases the pLDDT of the target IDR, and is used within a Metropolis acceptance criterion to either accept or reject such mutation. In this way, in practice, we favor mutations that increase the pLDDT of the target IDR. We iterate this algorithm until convergence, arriving at the final designed sequence. As a proof of principle, we apply the protocol to four different interesting targets: α-synuclein (ASYN), CD28, p53 (P53) and one of the small ubiquitin-like modifiers (SUMO). Possibly due to the lack of clear binding pockets, these proteins have been difficult to target using small-molecule drugs. Here, we selected these proteins due to the presence of an IDR in their structure, which serves as the binding epitope in our design protocol, and because they regulate biological processes whose disruption has clear downstream pathological consequences, making them compelling therapeutic targets. For example, ASYN has been shown to aggregate into toxic species that are implicated in the pathogenesis of Parkinson’s disease.^17^ CD28 can provide a co-stimulatory signal necessary for T-cell activation and survival, making it relevant in modulating immune responses in various diseases.^18^ The tumor suppressor protein (P53) can have its function restored by inactivating its interaction with negative regulators like MDM2, prevalent in many cancers, resulting in the inhibition of tumor growth.^19^ Lastly, SUMO proteins are an attractive drug target as SUMO’s modification of various substrates is involved in key cellular processes, while dysregulation of SUMOylation has been linked to diseases including cancer.^20^

**Figure 1.**
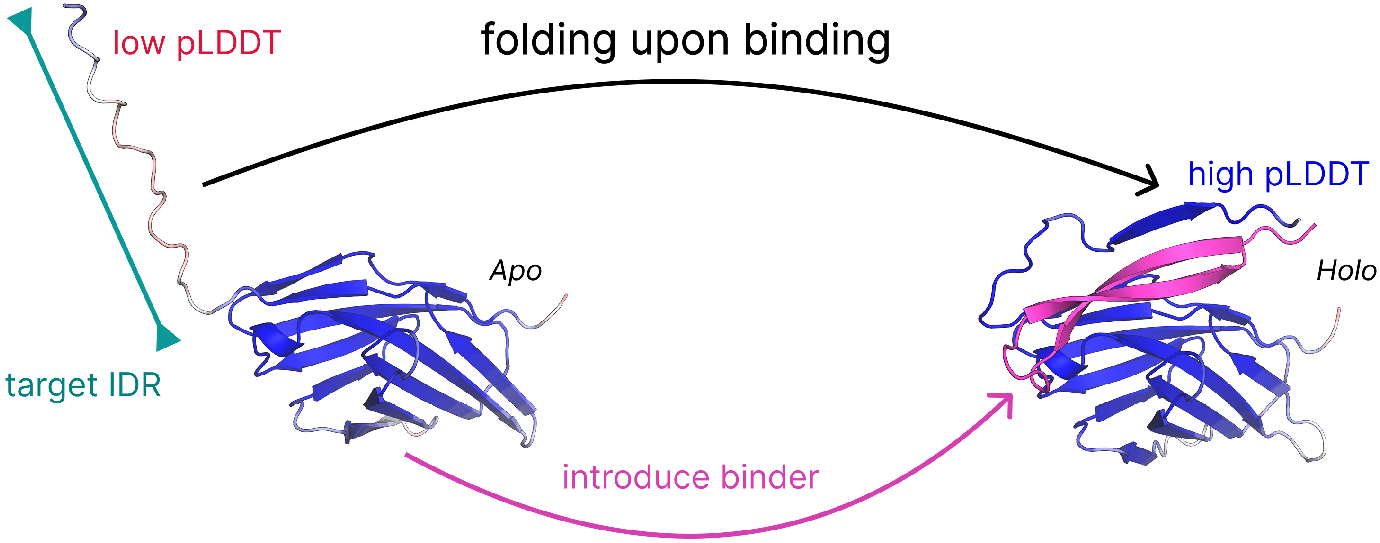
Schematic summary of our approach, showcasing inducing secondary structure and high model confidence upon the introduction of a binder (red-to-white-to-blue color represents low-to-high pLDDT values per residue). A protein target in its *apo* (unbound) form on the left has a disordered, unfolded region (white-to-red color). In the *holo* (bound) form on the right, upon interacting with a designed sequence (magenta), the previously unstructured region is induced to fold. The change in structuring is measured by a jump in the local pLDDT of the target region. A (positive) change in this local pLDDT is the main driving force during our Monte Carlo optimization in search of a candidate binder sequence.

Figure 2 compares the experimentally determined structures of the targets from the Protein Data Bank^21^ with the structures predicted by ESMFold for the isolated proteins. With the exception of ASYN, which is fully disordered^22^ and exhibits a high *C*_*α*_ Root Mean Square Deviation (RMSD) of 18.2Å and a low Template Modeling Score (TM-score)^23^ of 0.210, ESMFold accurately reconstructs the structured, non-IDR regions of CD28, P53, and SUMO, yielding low RMSDs (2.06, 2.12, and 0.683Å) and high TM-scores (0.762, 0.917, and 0.719), respectively. Aside from ASYN, the pLDDT is consistently high for the ordered regions (pLDDT-OR in Fig. 2), while being notably lower for their respective IDRs (pLDDT-IDR), reflecting reduced model confidence in the disordered parts. Importantly, for CD28, P53, and SUMO, the IDRs are absent in the experimental structures due to crystallographic challenges in resolving flexible regions. The selected targets exhibit varying degrees of disorder: ASYN lacks stable tertiary structure but exhibits transient secondary-structure propensity;^24^ CD28 contains a well-defined IDR; P53 has two disordered regions, both of which are selected as epitope targets; and SUMO contains two IDRs, of which we target only one. Molecular dynamics simulations further support the disordered nature of these regions, showing reduced intrinsic secondary structure content, while exhibiting elevated thermal fluctuations (see the Supplementary Information). Together, these observations strongly suggest that low model confidence (low pLDDT) for these systems correlates with intrinsic disorder, rather than reflecting the protein-folding algorithm’s failure to predict a well-structured region.

**Figure 2.**
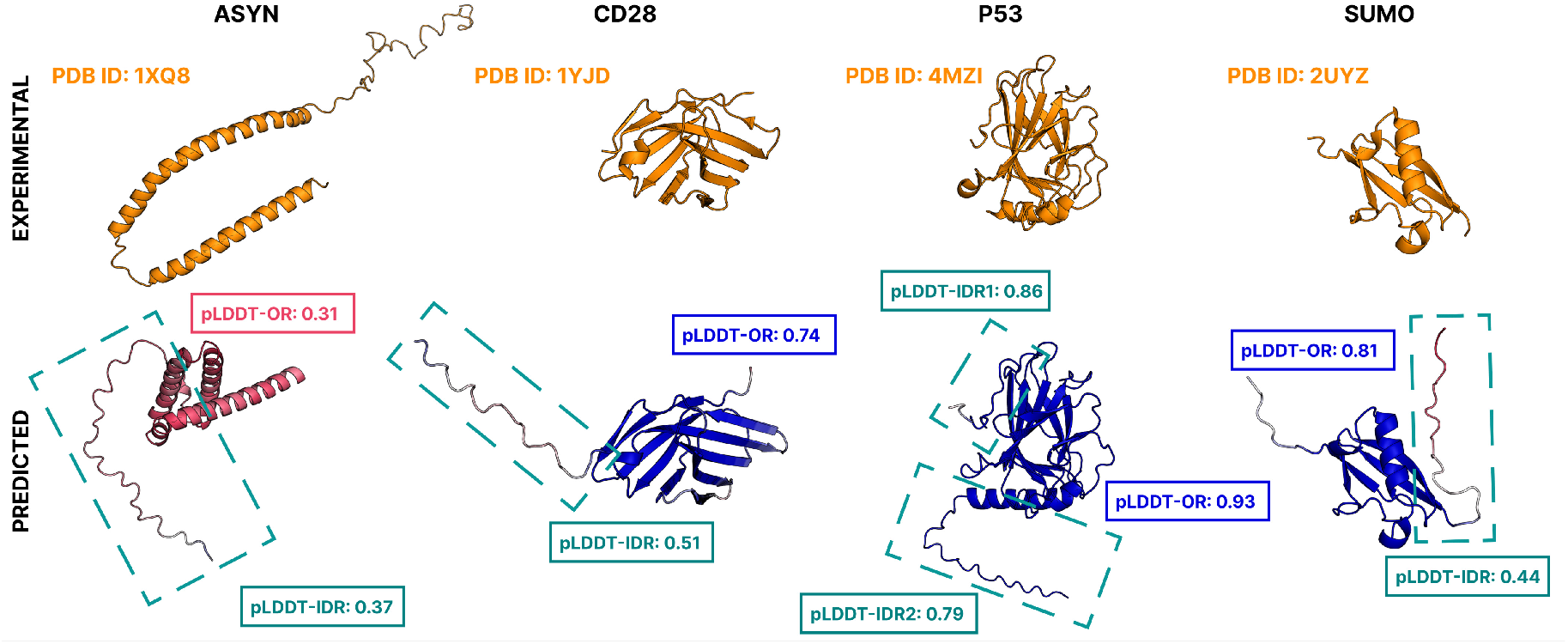
Comparison of experimental (top row, orange) and predicted (bottom row) structures by ESMFold for the four targets selected in this study. Predicted structures are colored per residue by pLDDT, using a red-to-white-to-blue gradient corresponding to values from 0 to 1. We report the average pLDDT over the ordered, rigid regions (pLDDT-OR) and the intrinsically disordered regions (pLDDT-IDR). Dashed teal boxes highlight the IDRs of interest (we omit SUMO’s second IDR for clarity). ESMFold accurately reproduces the *apo* form for all proteins except the fully disordered ASYN. IDRs forming unstructured coils are either unresolved in experimental structures (CD28, P53, SUMO), aligned well with experiment (P53), or poorly correlated (ASYN). For P53, we consider both termini (IDR1, IDR2) for epitope targeting. Although ESMFold predicts IDR1 with moderately high pLDDT, its coil-like geometry and enhanced thermal fluctuations (see the Supplementary Information) indicate a disordered character. On average, pLDDT-IDR is lower than pLDDT-OR (excluding ASYN), supporting the interpretation that ESMFold’s low confidence reflects intrinsic disorder rather than poor predictive accuracy. These disordered regions are subsequently used as epitope targets.

### B. ESMFold associates co-folding with an increase in pLDDT

Using our Monte Carlo optimization, we designed 11 binders across the four different targets previously described. Figure 3 presents the predicted structure and the per-residue pLDDT for targets in an isolated (unbound) form and in the presence of the binders designed using our protocol, highlighting the change in pLDDT upon binding. Both in the structure and in the pLDDT histogram, the region of the protein presented in teal corresponds to the region targeted during the design procedure. It is clear that, in general, upon binding the confidence level in the structural prediction increases for the targeted region. Apart from P53-1, where an increase in confidence level does not change the character of the secondary structure of the IDR, which remains a random coil, in all other targets presented here the change in pLDDT is also associated with a change in secondary structure, with the partial formation of alpha-helices or beta-sheets in the targeted region. While our procedure allows us to find binders for all targets, they also show different behaviors. The low pLDDT of ASYN, suggesting no stable tertiary structure despite some secondary structure propensity, is significantly increased in the presence of the binder, potentially correlating with an increased likelihood of obtaining a single dominant tertiary structure. In the case of CD28 and SUMO instead, the positive change in pLDDT is very much confined to the targeted IDR only (residues Gly121–Arg140 for CD28 and residues Gly1–Gly21 for SUMO). Finally, P53 represents yet another different situation, where the binder successfully induces a higher pLDDT region in residues Pro1–Val5, part of the targeted region, but at the same time decreases the pLDDT score in another, separate IDR (residues His200–His226). While this effect shows that our binders only induce structuring in their putative target region, it is unclear whether the change to lower values in other regions is a genuine physical mechanism of induced disorder (which we do not investigate further here), or from limitations in the pLDDT metric itself. As previously discussed, pLDDT can conflate model confidence with structural order. The presence of the binder might require ESMFold to attempt to predict complexes from sequence space farther away from its training data, simply lowering its confidence, particularly in regions that the model was already relatively less confident in even in the isolated state (as in the case of P53). Before we continue, we notice that while for ease of visualization we report here the results for a single binder per protein only, these sequences were chosen purely as representative examples. The very same trends are observed across multiple designs (see the Supplementary Information), confirming the robustness of our arguments and the design protocol. There we also show that for P53, we designed P53-E that targets the other IDR epitope, highlighting the flexibility of our method.

**Figure 3.**
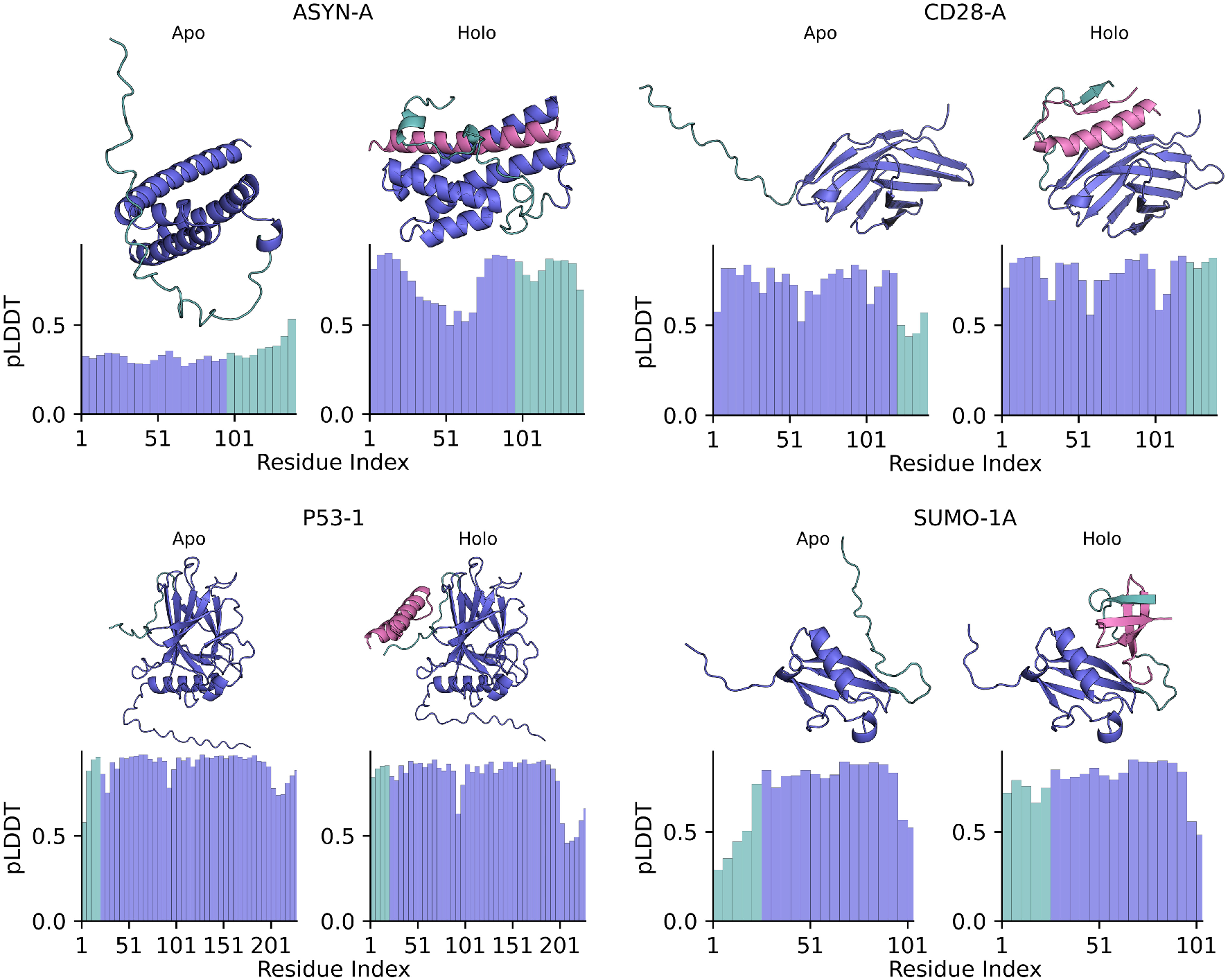
Four examples of peptide binder (magenta) designs. The targeted IDRs are colored in teal. The bottom bar plots for each structure show the pLDDT as an average over 5 contiguous residues (blue for the whole structure except the targeted IDR, in teal). Remaining designs are provided in Figure 7 in the Supplementary Information.

### C. Molecular simulations confirm high-affinity binding to the disordered regions

In Figure 4, we show the free energy profile as a function of distance from the binding site for all our different binders, together with the estimated binding affinity Δ*G*. We computed these free energy profiles using coarse-grained molecular dynamics (MD) in combination with umbrella sampling and replica exchange, see Methods for details. Compared to experimental techniques, molecular simulations provide us with direct mechanistic insights about the binding modality. For example, besides measuring binding affinity, we also obtain a clear microscopic picture of the binder-epitope interface. We use as a simple yet informative collective variable (CV) to describe the binding reaction: the distance *r* between the center of mass (COM) of the binder and that of the *rigid*, structured region of the target. This region is defined by first considering all close contacts of the binder to the target, and then discarding all the ones that involve the residues on the IDR. Fig. 4 shows clear deep minima at low *r*, of tens of *k*_*B*_*T*, for all target-binder systems. The variability in the width of this minimum across the different target-binder pairs can be explained by the fact that the epitope is part of an IDR, whose flexibility allows it to move away from the COM of the rigid part of the protein, while still being bound to the binder. This effect can then be different for each target-binder pair. In Figure 5, we display representative snapshots from the MD trajectories, clearly showing the unbinding process at larger CV values and supporting our interpretation of the shape of the free energy profiles. Moreover, we provide the reweighted thermal fluctuations (RMSF) analysis as a function of the distance in Figure 8 in the Supplementary Information, showing reduced RMSF values for the IDR in the bound region of the CV space.

**Figure 4.**
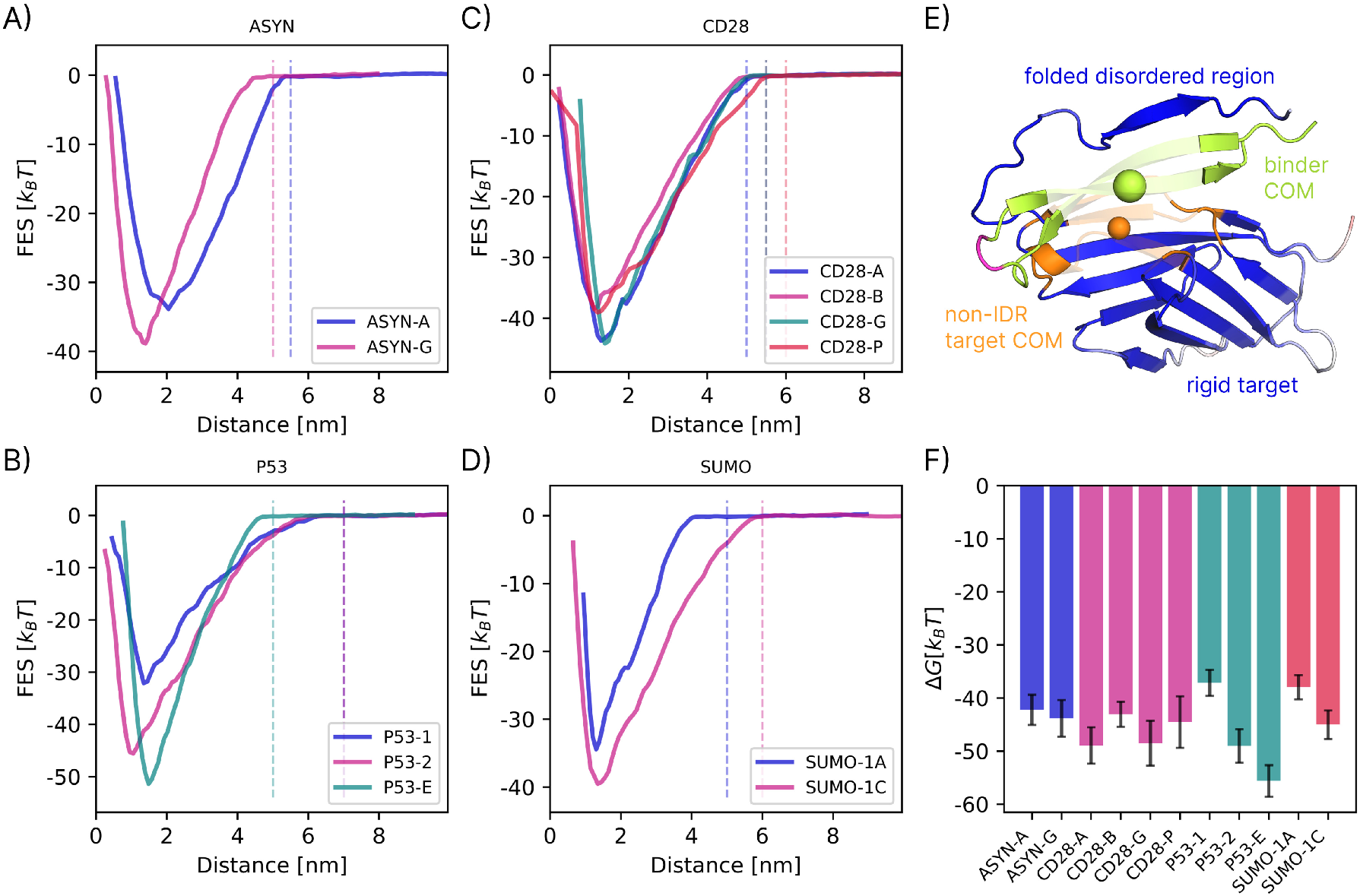
A–D) Free energy profiles along the distance *r*, computed using umbrella sampling with replica exchange. The dashed vertical lines represent the cutoff *r*_*c*_. E) Definition of the collective variable used in the free energy calculations. We measure the free energy as a function of the distance *r* between the centers of mass – depicted by the spheres – of the binder (lemon) and non-IDR part of the target (orange). F) Binding affinities (Δ*G*) for each system computed from the free energy surface (FES) projections in A–D) using Eq. 2, grouped in color by the protein target. Uncertainty is computed with block analysis,^25^ i.e., standard deviation of Δ*G* across 20 blocks.

**Figure 5.**
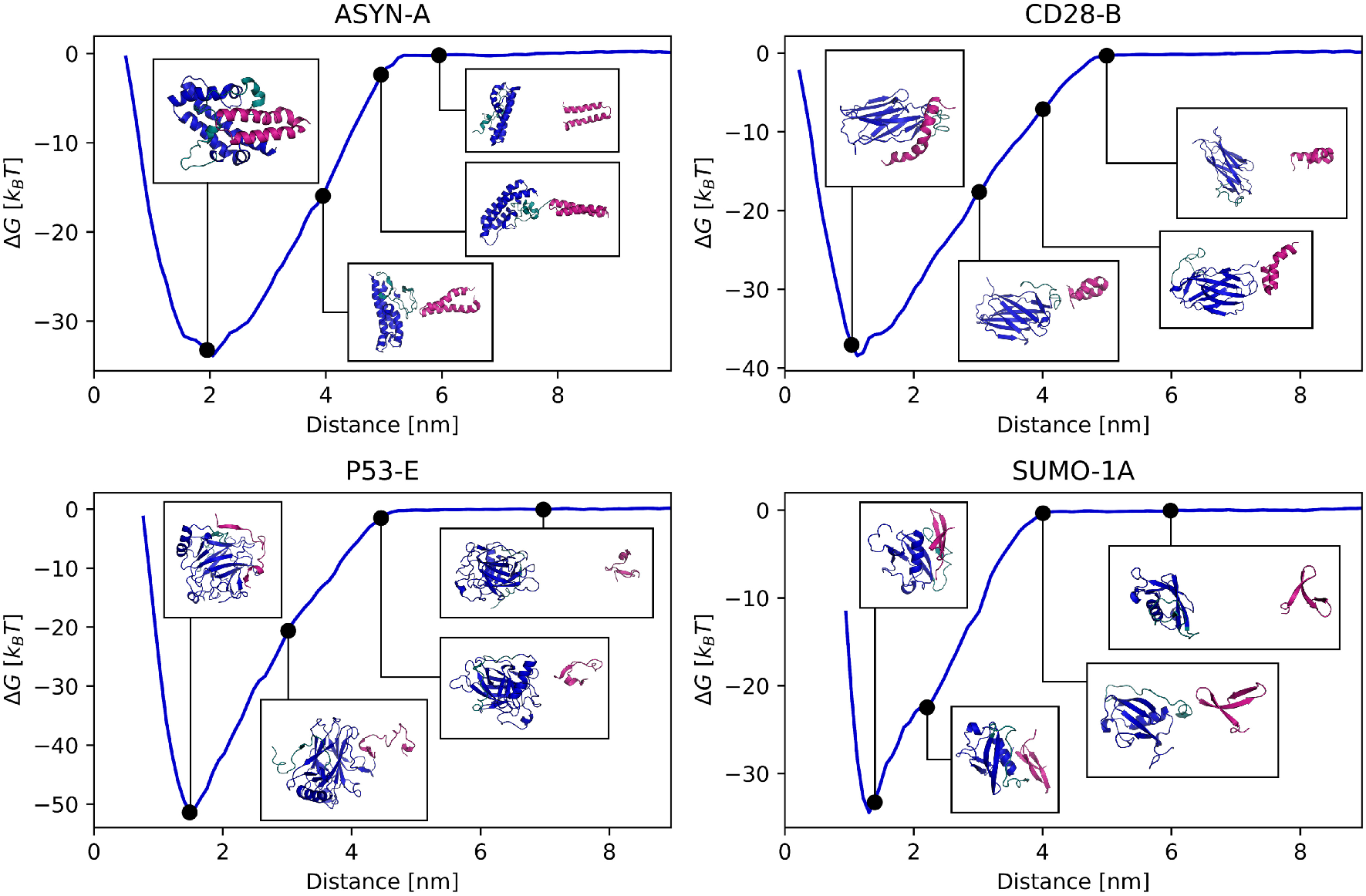
Four examples of binder-target systems together with snapshots from the trajectories at specific restraint distances *r*_*t*_ shown on the free energy surface (FES) as a black dot, with the visualization as an inset. At larger distances, the binder (magenta) is restrained away from the target (blue). The teal residues of the target show the disordered region.

An important aspect to highlight is that the affinity we obtain is typically associated with very strong binders such as antigen-antibody pairs. We expect these simulations to reproduce experimental binding measurements, such as surface plasmon resonance (SPR) or isothermal titration calorimetry (ITC), only within a few *k*_*B*_*T*, i.e., two or even three orders of magnitude in the dissociation constant *K*_*D*_. While various approximations in the coarse-grained description (e.g., the use of an implicit solvent) might have over-stabilized binding, the affinities we observe are large enough that even an over-estimation of many *k*_*B*_*T* would still strongly suggest that our sequences represent true, high-affinity binders. Additionally, it should be pointed out that MD simulations (more precisely, free energy calculations using MD), based on a description of the physico-chemical forces in the system, represent an independent, orthogonal approach compared to a deep learning-based protein-folding algorithm, which indirectly extracts information from existing databases of sequences and experimental structures. That two independent methods, i.e., ESMFold and MD, predict binding of our designed sequences to the target epitope strengthens our belief that these results will stand experimental validation, which, however, remains outside the scope of this work.

### D. Discussion

We have shown proof-of-concept of a design protocol for peptide binders targeting disordered regions on clinically important protein targets. Our protocol is based on the definition of an energy function, which can be evaluated from the structure predicted by a protein-folding algorithm (here, ESMFold), and its minimization as a function of the binder sequence, using a Monte Carlo approach. An important ingredient of such a function is the confidence metric associated with the intrinsically disordered region of the protein we target, whose increase we hypothesized is a signal for induced folding. Free energy calculations via molecular dynamics with a classical force field widely used for proteins confirm the *bona fide* nature of our designed binders, as well as strong binding specifically to their putative target epitope. In particular, our *in silico* validation shows promising binding affinities, providing good candidates that could be experimentally tested in a lab. In spirit, the approach we presented here is similar to what was recently shown by Baker and co-workers.^26^ These authors first designed a template library of peptide fragments that recognize specific repeating amino acid patterns, and then stitched these fragments into small proteins that could recognize extended patterns of disordered sequences. These small proteins were shown to bind strongly and selectively to their targets, with low cross-reactivity, and various proof-of-principle downstream applications were provided, from cell lysate separation to inhibition of cancer-associated proteins. Similarly to our work, their work showed that targeting of IDR is possible, our approach differs mostly for its simplicity: instead of a multi-step protocol with different models and several human interventions between different steps, we rely on a single-step design purely driven by an energy function directly evaluated from the output of a single protein-folding algorithm. We also notice that the binders we generated with our protocol, of only 30–50 residues, are much smaller than those designed in the aforementioned publication.^26^ The smaller size could provide translational advantages. First, it allows obtaining large quantities of such binders at high purity using solid-phase peptide synthesis techniques. Second, and perhaps more importantly, shorter lengths are associated with weaker interactions with other proteins and generally make these binders harder to recognize by the immune system, decreasing their immunogenic potential. Overall, having the ability to design small peptide binders on demand for disordered regions of proteins, previously considered untargetable via typical approaches, opens up a range of possibilities across the field of applied biotechnology and drug design.

## III. METHODS

### A. Binder-design protocol

The practical implementation of our protocol builds on Monte Carlo optimization, as implemented using the open-source `BAGEL` package.^27^ While an in-depth description of the protocol can be read in the associated paper,^27^ we describe here the general idea for completeness. The starting point is the definition of an energy (loss) function that depends on structural constraints imposed on different parts of the protein-binder complex. In particular, let *X* ∈ *χ* denote a candidate folded structure (i.e., the 3D atomic coordinates of the complex) and Φ ∈ **Φ** the associated folding-confidence metrics – e.g., pLDDT, pTM, Predicted Aligned Error (PAE) – predicted by a structure prediction oracle. We then define the total energy *E* : *χ* × Φ **→** ℝ as a weighted sum of individual energy (loss) terms, each of which is a function of either the structure or its confidence metrics:

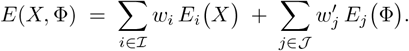

Here, *E*_*i*_(*X*) captures geometric or inter-residue constraints acting directly on the coordinates *X, E*_*j*_(Φ) captures folding-confidence penalties or rewards based on the metrics Φ, and *w*_*i*_, *w*_*j*_ control their relative importance respectively. In our implementation, we use a total of eight distinct terms (i.e., | ℐ | + | 𝒥 | = 8), whose different combinations produce various designs (detailed in the Supplementary Information). Besides terms previously reported for general protein design purposes, e.g., steering the design toward high confidence structures – high pLDDT, high pTM, low PAE – and soluble sequences (important to aid expression and experimental verification), our loss function contains additional custom terms that drive the designed binder to bind the selected IDR (rather than generically the protein), while also forcing the latter to fold upon binding. A detailed description of each term and associated weights *w* is reported in Table I in the Supplementary Information.

**Table I.** Energy terms of the protein design guided optimization of a binder targeting an IDR. First set of energy terms relates to structure-related properties *E*_*i*_(*X*), while the second set shows the folding-metric associated terms *E*_*j*_ (Φ). For detailed implementation, refer to the BAGEL package and technical report.^27^.

| Energy Terms | Description |
| --- | --- |
| HydrophobicEnergy | Reduces the number of hydrophobic residues (on the surface). |
| AverageDistance | Penalizes large distances between binder and the IDR. |
| SecondaryStructureEnergy | Promotes formation of a secondary structure (alpha-helices, beta-sheets) on the IDR. |
| PTMEnergy | Guides the complex toward high-confidence structures. |
| PAEEnergy | Minimizes interface PAE, i.e., having the binder and IDR behave as a single rigid body. |
| OverallPLDDTEnergy | Encourages the entire complex to be a highly confident prediction by the model. |
| PLDDTEnergy (binder) | Enforces additional confidence in the *de novo* designed binder. |
| PLDDTEnergy (epitope) | Enforces further ordering and rigidification of specific regions within the IDR. |

Once the energy function is defined, we search for the binder sequence by minimizing such energy through an adaptation of a Monte Carlo tempering algorithm. Even for short amino acid sequences, the number of potential peptide candidates is immense. Whereas this large selection increases the possibility that at least one candidate with the right properties exists, it is also an impossibly large space to search without any type of guidance. We solve this issue by using the energy function to drive the search process by a slightly modified version of a classical Monte Carlo algorithm called simulated tempering,^28^ to help sample the sequence space in search of the optimal (global) solution. We start from a random sequence, and at each iteration generate a new potential candidate via a random point mutation, choosing one of the 20 canonical amino acids with equal probability, except for cysteine (because of problems associated with expression and thus related to an eventual experimental validation). A mutation is accepted or rejected based on the Metropolis criterion, and this protocol is iterated until a maximum number of steps is reached, on the order of 10^4^ steps. If *s* ∈ 𝒮 is the current protein sequence, and *s*^′^ is a candidate mutated sequence, the acceptance probability is given by

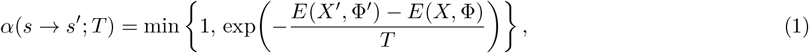

where (*X*, Φ) = ℱ (*s*) and (*X*^′^, Φ^′^) = ℱ (*s*^′^) are the folded structures and folding confidence metrics corresponding to sequences *s* and *s*^′^ respectively. In Monte Carlo tempering, the system cycles between a low temperature (*T* = 0.1) to promote exploitation and a high temperature (*T* = 1.0) to encourage exploration, using the acceptance probability defined in Eq. 1. Since we are interested in optimization rather than sampling of the energy landscape according to a Boltzmann distribution, instead of trying to swap between the two temperatures using an appropriate acceptance criterion to maintain detailed balance, we simply cycle between the low and high temperature regimes sequentially (usually every *n*_low_ = 400 and *n*_high_ = 100 steps, although the exact values are system-dependent). At the end of each high temperature regime, we also switch back to the system (i.e., the sequence) with the lowest energy sampled so far. In general, while in the low temperature regime the system efficiently explores its current local minimum, at high temperature, the system can jump between minima separated by larger energy barriers more easily, providing an efficient balance between local and global exploration. While other schemes could be used, such as parallel tempering,^29^ a comprehensive benchmark is beyond the scope of this work. We report the values of the different energy terms for the 11 binders we designed in Table VI in the Supplementary Information. We notice that the final values obtained for the different energy terms obtained at the end of the optimization for our designs are consistent with high-quality binders, with interface PAE (iPAE) values around or below 0.2, (local) pLDDT scores around or above 0.75, and binder–epitope distances around or under 1.0 nm.

### B. Molecular simulations

#### 1. Simulation setup and force field

We describe the interactions with the Amber ff14SB force field^30^ and an implicit water model.^31^ To simulate the system, we used OpenMM^32^ with PLUMED.^33^ After fixing the final structures of the Monte Carlo optimization from ESMFold with `pdbfixer`,^32^ we minimized the bound complex with L-BFGS in two phases: first, we restrained the non-hydrogen atoms and minimized the rest of the complex; second, we minimized all atoms together. We ran 10 ns of equilibration in the NVT ensemble, extracting the final structures as the starting point for our free energy calculations. Given we used an implicit solvent, we did not use periodic boundary conditions, while non-bonded interactions were cut off at a distance of 2 nm, with the solute dielectric set to 1.0 and the solvent dielectric to 78.5 (dimensionless relative permittivities). All simulations are performed with a timestep of 2 fs.

We recognize that using an implicit solvent and cutoff electrostatics are suboptimal compared to describing the energetics of the system using an explicit water model with periodic boundary conditions and a proper treatment of long-range effects using schemes such as Particle Mesh Ewald.^34^ However, we chose our simpler setup to at least ensure accurate statistical sampling and converged free energy profiles, which would instead be problematic with more computationally expensive (but still inaccurate) models. While we initially explored more accurate energetic descriptions, including explicit solvent simulations with periodic boundary conditions, the presence of highly flexible IDRs would have required prohibitively large simulation boxes - a limitation avoided by using an implicit solvent model. Although using an implicit solvent model could lead to an inaccuracy of several *k*_*B*_*T* in the calculated binding free energy, our aim is to qualitatively confirm the presence of a deep minimum associated with the bound state, and that this minimum is associated with a formation of a folded structure. Given the values of several *tens* of *k*_*B*_*T* we obtain for our minima, we believe these predictions to be qualitatively robust to changes in the simulation setup.

#### 2. Free energy calculations and binding affinity estimation

To validate our results, in particular, to verify that the designed sequences have appreciable binding to their target, we employed free energy calculations using a combination of umbrella sampling^35^ and replica exchange. The former is employed to obtain the free energy as a function of the distance between the center of mass (COM) of the target and the binder, since its integral in the bound state can be used to gauge the strength of binding, and is formally related to the experimental binding/dissociation constants.^36^ More precisely, one can estimate the free energy of binding Δ*G* for an observed one-dimensional CV free energy profile *F*_obs_(*r*) with the formula

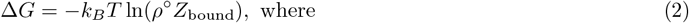

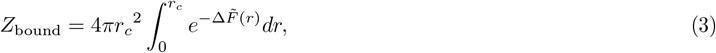

where 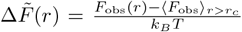, *k*_*B*_ is the Boltzmann’s constant, *ρ*° is the standard reference concentration, i.e., 1 mol L^−1^, and *r*_*c*_ is the cutoff distance defining the boundary between the bound and unbound states. 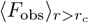 is the effective estimate of *F* (∞), where 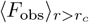 is the mean of the free energy averaged between its values recorded between *r*_*c*_ and max({*r*_*t*_}). The boundary *r*_*c*_ is defined as the distance above which the free energy *F* (*r*) remains constant, and thus is equal to its value at infinite distance, where the protein-binder pairs do not interact. We introduced the average to remove the effect of random fluctuations in the estimates of this plateau. A full derivation of the above formula and an in-depth discussion of the different terms is provided in the Supplementary Information. While doing umbrella sampling, we bias the system using as a collective variable (CV) the distance *r* between the COMs, *x*_target_ and *x*_binder_, of groups of *C*_*α*_ atoms for residues on the target and binder respectively. We defined the group on the target by finding the nearby residues close to the binder after the initial equilibration. We excluded the residues that form a part of the targeted IDR epitope, as otherwise the large fluctuations of the IDR made *x*_target_ lie outside of the rigid domain of the target, leading to degeneracies in *r*. With this definition, the bias used during umbrella sampling is

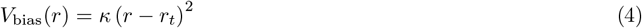

which forces the system to sample close to the region *r* ≈ *r*_*t*_, allowing one to obtain an accurate value for the local free energy. As customary in umbrella sampling, different simulations (windows) are repeated with biases differing in the value of *r*_*t*_, and then stitched together using the Weighted Histogram Analysis Method (WHAM) algorithm^37^ to obtain the free energy profile. The exact values of all the parameters used in our simulations, including the values of the different bias centers {*r*_*t*_} or the bound region cutoffs *r*_*c*_, can be found in the Supplementary Information.

To further improve sampling, especially of the other slow degrees of freedom involved in the binding transition, e.g., torsional angles describing the state of the IDR and the binder, we employed replica exchange for each window. In particular, for each value of the external bias used in umbrella sampling, we run four replicas in parallel using OpenMMTools.^38^ The replicas have different temperatures, separated by 5 kelvin each, ranging from 310 K to 325 K. During the simulations, we attempt *n*^3^ swaps between two replicas chosen randomly every 10 ps, where *n* is the number of replicas. Overall, after initial 10 ns of equilibration within the replica exchange setting, all systems were simulated for 30 ns for each (bias, replica temperature) combination, which amounts to about 10 µs of total simulation time per protein-binder pair. Details of acceptance rates and sampling convergence can be found in the Supplementary Information.

## C. Data, Materials, and Software Availability

All data needed to evaluate the conclusions in the paper are present in the paper and/or the Supplementary Information. We used the open-source Python package `BAGEL`^27^ available athttps://github.com/softnanolab/bagel, with version v0.1.0 archived in the Zenodo repository.^39^ The templates to obtain the binders can be found in the GitHub and Zenodo repositories in the folder `scripts/technical-report/disordered/`.

## Acknowledgments

We thank Dr Daniele Visco for insightful discussions about the problem and implementation of some of the simulation techniques used. J.L. acknowledges the President’s PhD scholarship at Imperial College London for funding. We acknowledge computational resources and support provided by the Imperial College Research Computing Service (http://doi.org/10.14469/hpc/2232). We are grateful to the UK Materials and Molecular Modelling Hub for computational resources, which is partially funded by EPSRC (EP/T022213/1, EP/W032260/1 and EP/P020194/1). This research was also supported by grants from NVIDIA and utilized NVIDIA’s GPU access through the Academic Grant for Simulation and Modeling.

## E. Author contributions

S.A.-U. designed the research; J.L. and S.A.-U performed the research, analyzed the data and wrote the manuscript.

## SUPPLEMENTARY INFORMATION

### 1. Analysis of IDR secondary structure and fluctuations

To compute the TM-score of the binder-target complex as predicted by ESMFold, we use the `tmalign` PyMOL implementation of the `TM-Align` package.^40^ For RMSD calculation we use `biotite`,^41^ and for Root Mean Square Fluctuations (RMSF)^42^ and secondary structure assignment (through DSSP^43^) we use `MDAnalysis`^44^.^45^

Figure 6 shows the thermal fluctuations of the disordered regions of interest, highlighted in teal, including the RMSF and dominant secondary structure per residue. For the per-residue RMSF computation, we use the average *C*_*α*_ positions aligned to the non-IDR parts only. For P53, we focus on two different IDRs to showcase the flexibility of our design protocol - one of which is clearly more disordered (with a high RMSF, on the right), while the other has a relatively moderate RMSF, but contains coil-like characteristics (on the left).

**Figure 6.**
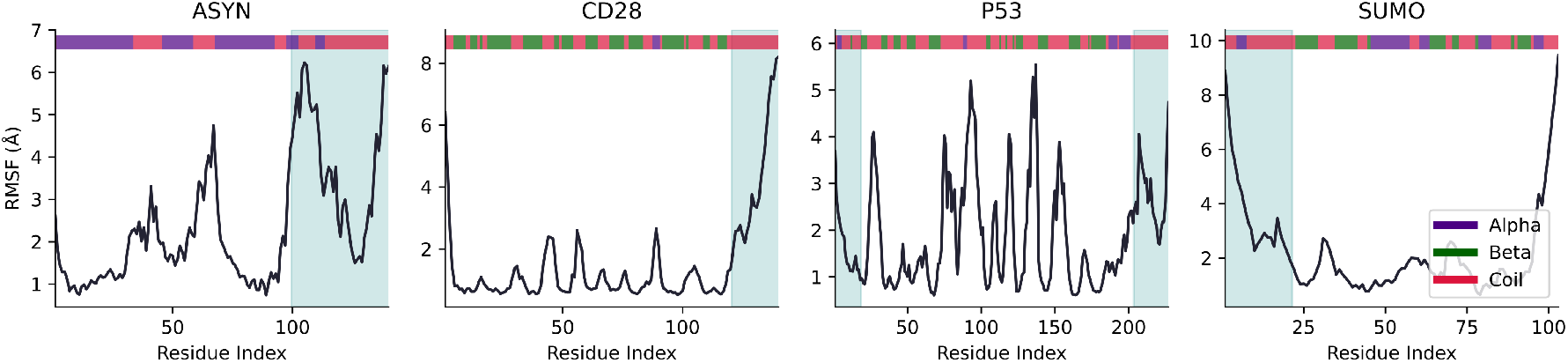
Molecular dynamics of 500 ns of the four targets at 310 K in the *apo* form, using otherwise the same simulation parameters (force field, etc.) as the enhanced sampling simulations in the free energy calculations, but without the harmonic restraint, nor the replica exchange. IDRs (teal shade) of the protein have elevated *C*_*α*_ RMSF and have generally greater coil-like nature in their secondary structure compared to the rest of the protein.

### 2. Details of Monte Carlo optimization

Table I describes all energy terms employed in the protocol. Tables II to V then provide the specific energy terms, *E*_*i*_ and *E*_*j*_, and their weights, *w*_*i*_ and *w*_*j*_ respectively, used for the different design protocols, including the details of the optimization. The exact weights need to be carefully considered and tuned to achieve the desired behavior. We have conceived of the terms and their weights through rational design and trial-and-error. Note that given the stochastic nature of the optimization, using these optimization parameters will not yield the exact binders as discussed here. For full description of these energy terms, consult the `BAGEL` package and its associated technical report.^27^

**Table II.** Peptide binders for ASYN, specifically P37840 (SYUA HUMAN). Hotspot residues are Leu100–Ala140. All binders have a length of 50 residues.

|  | ASYN-A | ASYN-G |
| --- | --- | --- |
| Energy Terms | Weights $w$ | |
| PTMEnergy | 1.0 | 1.0 |
| OverallPLDDTEnergy | 1.0 | 1.0 |
| HydrophobicEnergy | 2.0 | 2.0 |
| PAEEnergy | 4.0 | 4.0 |
| AverageDistance | 0.2 | 0.2 |
| PLDDTEnergy (binder) | 4.0 | 4.0 |
| PLDDTEnergy (epitope) | — | 4.0 |
| SecondaryStructureEnergy (α-helix) | 4.0 | — |
| Optimization | Metric |  |
| $T_{\text{low}}$ | 0.2 | 0.2 |
| $T_{\text{high}}$ | 2.0 | 2.0 |
| $n_{\text{high, steps}}$ | 50 | 50 |
| $n_{\text{low, steps}}$ | 50 | 50 |

**Table III.** Peptide binders for CD28, specifically P10747 (CD28 HUMAN). Hotspot residues Gly121–Arg140. All binders have a length of 30 residues.

|  | CD28-A | CD28-B | CD28-G | CD28-P |
| --- | --- | --- | --- | --- |
| **Energy Terms** | **Weights $w$** |  |  |  |
| PTMEnergy | 1.0 | 1.0 | 1.0 | 1.0 |
| OverallPLDDTEnergy | 1.0 | 1.0 | 1.0 | 1.0 |
| HydrophobicEnergy | 2.0 | 2.0 | 2.0 | 2.0 |
| PAEEnergy | 4.0 | 4.0 | 4.0 | 4.0 |
| AverageDistance | 0.2 | 0.2 | 0.2 | 0.2 |
| PLDDTEnergy (binder) | 4.0 | 4.0 | 4.0 | 4.0 |
| PLDDTEnergy (epitope) | 4.0 | 4.0 | 4.0 | — |
| SecondaryStructureEnergy (α-helix) | 8.0 | — | — | — |
| SecondaryStructureEnergy (β-sheet) | — | 8.0 | — | — |
| **Optimization** | **Metric** |  |  |  |
| $T_{\text{low}}$ | 0.3 | 0.1 | 0.1 | 0.1 |
| $T_{\text{high}}$ | 2.0 | 1.0 | 1.0 | 1.0 |
| $n_{\text{high, steps}}$ | 100 | 100 | 100 | 100 |
| $n_{\text{low, steps}}$ | 100 | 100 | 100 | 100 |

### 3. Further results for the optimal binders

Table VI reports the most relevant metrics typically used to analyze protein-protein complexes and their binding interface, for the 11 final binders designed, in the bound state with their target. As observable from the weights of the different energy terms involved in the optimization, as reported in Tables II to V, the energy terms associated with iPAE, and the local pLDDT (over the binder and over the epitope) are the key metrics driving our minimization protocol. Finally, Figure 7 provides the remaining designs and pLDDT analysis (as in Figure 3 in the main text), for the binders not shown in the main body of the text.

**Table IV.** Peptide binders for P53, specifically P04637 (P53 HUMAN). P53-1 and P53-2 target epitope residues Pro1–Phe18, while P53-E targets the other disordered epitope at His205–His227. All binders have a length of 30 residues.

|  | P53-1 | P53-2 | P53-E |
| --- | --- | --- | --- |
| **Energy Terms** | **Weights $w$** |  |  |
| PTMEnergy | 1.0 | 1.0 | 1.0 |
| OverallPLDDTEnergy | 1.0 | 1.0 | 1.0 |
| HydrophobicEnergy | 2.0 | 2.0 | 2.0 |
| PAEEnergy | 5.0 | 5.0 | 5.0 |
| AverageDistance | 0.2 | 0.2 | 0.2 |
| PLDDTEnergy (binder) | 5.0 | 5.0 | 5.0 |
| PLDDTEnergy (epitope) | 5.0 | 5.0 | 5.0 |
| **Optimization** | **Metric** |  |  |
| $T_{\text{low}}$ | 0.1 | 0.1 | 0.1 |
| $T_{\text{high}}$ | 1.0 | 1.0 | 1.0 |
| $n_{\text{high, steps}}$ | 200 | 200 | 200 |
| $n_{\text{low, steps}}$ | 200 | 200 | 200 |

**Table V.** Peptide binders for SUMO, specifically P63165 (SUMO1 HUMAN). Epitope residues are Gly1–Gly21. All binders have a length of 30 residues.

|  | SUMO-1A | SUMO-1C |
| --- | --- | --- |
| Energy Terms | Weights $w$ | |
| PTMEnergy | 1.0 | 1.0 |
| OverallPLDDTEnergy | 1.0 | 1.0 |
| HydrophobicEnergy | 2.0 | 2.0 |
| PAEEnergy | 5.0 | 5.0 |
| AverageDistance | 0.2 | — |
| PLDDTEnergy (binder) | 5.0 | — |
| PLDDTEnergy (epitope) | 5.0 | 5.0 |
| Optimization | Metric |  |
| $T_{\text{low}}$ | 0.1 | 0.1 |
| $T_{\text{high}}$ | 1.0 | 1.0 |
| $n_{\text{high, steps}}$ | 200 | 200 |
| $n_{\text{low, steps}}$ | 200 | 200 |

**Table VI.** Metrics for selected energy (loss) terms in the IDR-guided design protocol. We report the actual values for these (unweighted) rather than the (weighted) energy term values. iPAE is normalized.

| System | Separation [nm] | iPAE [a.u.] | Binder pLDDT [a.u.] | Epitope pLDDT [a.u.] |
| --- | --- | --- | --- | --- |
| ASYN-A | 0.796 | 0.12 | 0.87 | 0.81 |
| ASYN-G | 0.676 | 0.21 | 0.85 | 0.66 |
| CD28-A | 1.002 | 0.12 | 0.88 | 0.85 |
| CD28-B | 0.893 | 0.16 | 0.86 | 0.78 |
| CD28-G | 0.938 | 0.16 | 0.84 | 0.71 |
| CD28-P | 0.704 | 0.18 | 0.81 | 0.74 |
| P53-1 | 1.244 | 0.12 | 0.88 | 0.88 |
| P53-2 | 0.944 | 0.13 | 0.89 | 0.88 |
| P53-E | 2.402 | 0.31 | 0.88 | 0.68 |
| SUMO-1A | 0.538 | 0.20 | 0.77 | 0.69 |
| SUMO-1C | 0.960 | 0.16 | 0.87 | 0.82 |

**Figure 7.**
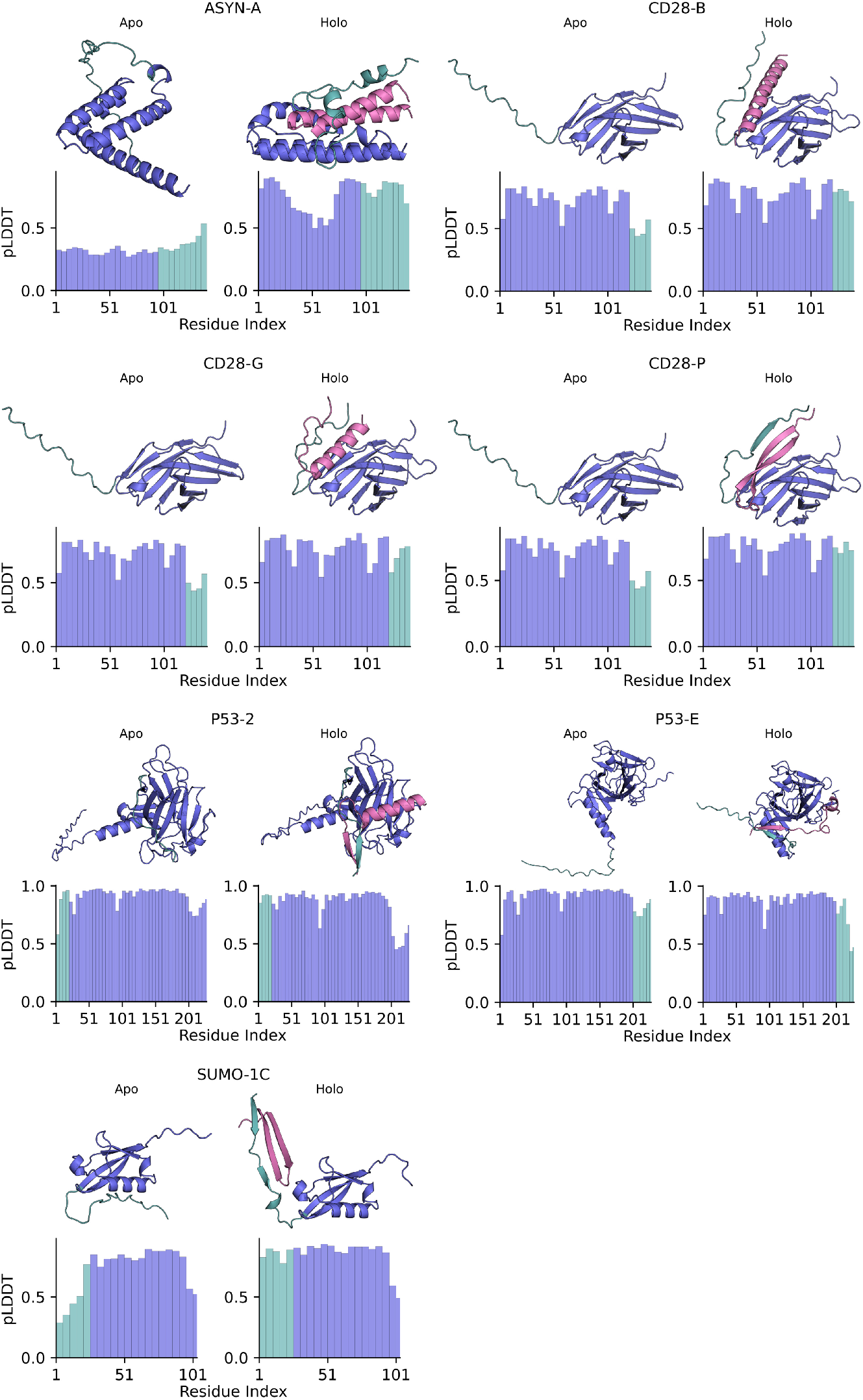
All remaining peptide binder (magenta) designs, targeting the epitope (teal) of the target (blue). The bottom bar plots for each structure also show the pLDDT as an average over 5 contiguous residues.

**Figure 8.**
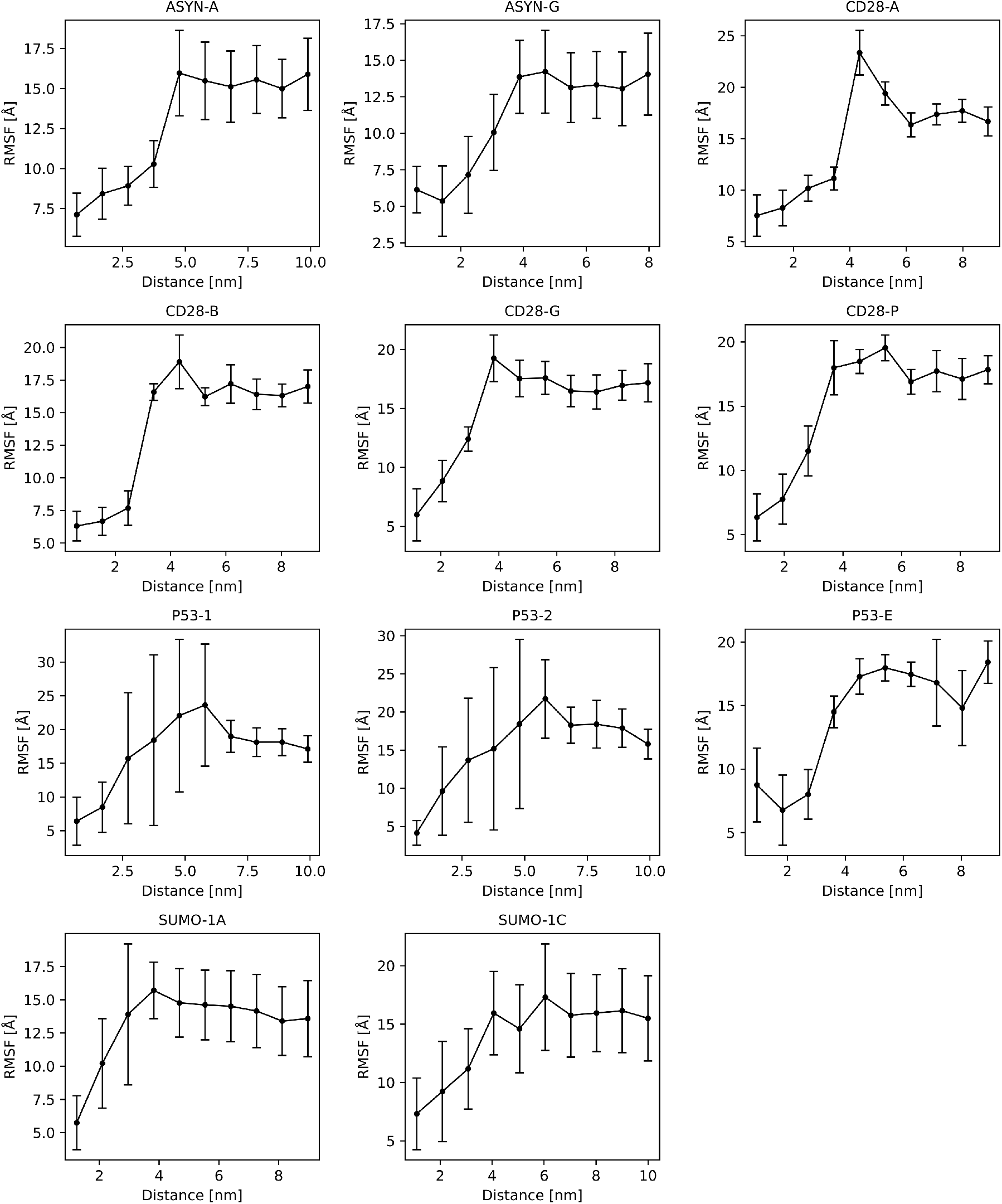
ℝoot Mean Square Fluctuations (RMSF) for all systems as a function of the CV distance. The CV space is divided into 10 equal-sized bins and reweighted according to the restraint bias. Each point here represents the mean RMSF across all residues of the IDR;, and the vertical bars show the standard deviation across those residues.

### 4. Simulation and analysis details for free energy calculations

#### a. Collective variable

We define the distance *r*(*x*_target_, *x*_binder_) between the centroids of two *C*_*α*_ atom groups, *X*_target_ and *X*_binder_. Here, *x*_target_ and *x*_binder_ denote the centroids of their respective groups. Each group comprises all residues whose *C*_*α*_ atoms lie within 1.5 nm of any *C*_*α*_ in the other group. To focus on ordered regions, any residues identified as part of an IDR – by visual inspection of the equilibrated structure – are excluded from *X*_target_. This was done in order to prevent artifacts introduced by large thermal fluctuations of the IDR, which would often lie outside of the rigid domain. Moreover, when *x*_target_ and *x*_binder_ overlap, there would be an inherent degeneracy, where different binding modes collapse into a single point on the free energy surface.

#### b. Umbrella sampling

Table VII provides the details of window placements for umbrella sampling. We chose these restraints based on a first of set of well-tempered metadynamics simulations^46^ not included here, which suggested the width of the bound basin in our CV space. In the presumed bound region of the CV space, we place windows at a higher density (narrow) compared to the unbound region with a lower density (wide). Starting configurations of the simulations were retrieved by displacing the binder along the vector **r** pointing from *x*_target_ to *x*_binder_ to the respective values of *r*_*t*_ for that specific window. Subsequent equilibration within the replica exchange setting ensured the initial conformation remains physical. If the binder displacement caused steric clashes, e.g., when displacing the binder closer to the target, the nearest *valid* non-sterically clashing starting configuration was used instead, where then the associated harmonic restraints alone are responsible for moving the binder to the local region of the CV space for that particular window within the equilibration phase.

**Table VII.** Umbrella sampling with replica exchange simulation windows. Narrow windows use *κ*_narrow_ = 412.40 kJ mol^−1^ nm^−2^ with 0.125 nm spacing and wide windows use *κ*_wide_ = 103.10 kJ mol^−1^ nm^−2^ with 0.250 nm spacing. The bound/unbound cutoff *r*_*c*_ is defined from visual inspection of the plateau in the free energy profiles and is used to compute Δ*G*.

| System | Narrow Windows | Wide Windows | $r_c$ [nm] |
| --- | --- | --- | --- |
| | $r_t$ [nm] | $r_t$ [nm] | |
| ASYN-A | 0.000, 0.125, ..., 7.750, 7.875 | 8.000, 8.250, ..., 9.750, 10.000 | 5.5 |
| ASYN-G | 0.000, 0.125, ..., 5.750, 5.875 | 6.000, 6.250, ..., 7.750, 8.000 | 5.0 |
| CD28-A | 0.000, 0.125, ..., 6.750, 6.875 | 7.000, 7.250, ..., 8.750, 9.000 | 5.0 |
| CD28-B | 0.000, 0.125, ..., 6.750, 6.875 | 7.000, 7.250, ..., 8.750, 9.000 | 5.5 |
| CD28-G | 0.000, 0.125, ..., 6.750, 6.875 | 7.000, 7.250, ..., 8.750, 9.000 | 5.5 |
| CD28-P | 0.000, 0.125, ..., 6.750, 6.875 | 7.000, 7.250, ..., 8.750, 9.000 | 6.0 |
| P53-1 | 0.000, 0.125, ..., 8.750, 8.875 | 9.000, 9.250, ..., 9.750, 10.000 | 7.0 |
| P53-2 | 0.000, 0.125, ..., 7.750, 7.875 | 8.000, 8.250, ..., 9.750, 10.000 | 7.0 |
| P53-E | 0.000, 0.125, ..., 6.750, 6.875 | 7.000, 7.250, ..., 8.750, 9.000 | 5.0 |
| SUMO-1A | 0.000, 0.125, ..., 6.750, 6.875 | 7.000, 7.250, ..., 8.750, 9.000 | 5.0 |
| SUMO-1C | 0.000, 0.125, ..., 7.750, 7.875 | 8.000, 8.250, ..., 9.750, 10.000 | 6.0 |

#### c. Extracting the binding free energy Δ*G* from the free energy profiles: full derivation

Here, we assume knowledge of textbook statistical mechanics and provide a semi-exhaustive derivation of Eq. 3, specifically considering all the corrections necessary for the free energy profile in a one-dimensional collective variable of a molecular system of 3*N* total coordinates, where *N* is the number of atoms in the system.

Let **r** be the vector connecting the centers of mass of the binder and the rigid part of the target, i.e., connecting groups *X*_target_ and *X*_binder_, with *r* = ∥**r**∥. The three–dimensional potential of mean force (PMF) *F*_true_(**r**) is defined by the probability distribution

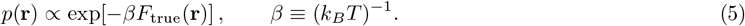

Notice that *F*_true_(**r**) is a free energy because it is itself the result of integrating over all the degrees of freedom of the protein and the binder, at a specific value of **r**. The standard–state binding free energy for a bimolecular association is:

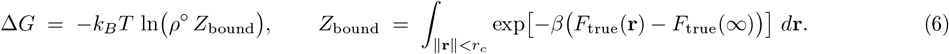

Rewriting the previous equation in spherical coordinates we have

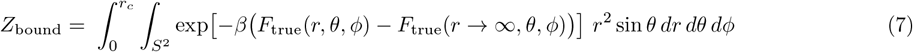

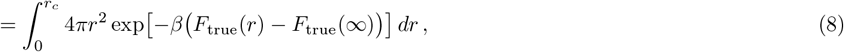

where we define

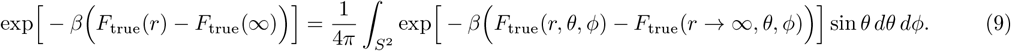

Here, the integral is over the solid angle *S*^2^ with a measure *d*Ω = sin *θ dθ dϕ*, where 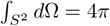.

In our simulations, we reconstruct a one–dimensional free energy profile *F*_obs_(*r*) with WHAM. This free energy differs from the radial PMF defined by Eq. 9 by the Jacobian of the change of variables r ≡ (*x, y, z*) → (*r, θ, ϕ*):

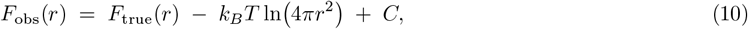

with *C* being an arbitrary additive constant. We refer to this as the *entropic contribution* arising from a larger density of states at larger *r*. Substituting Eq. (10) into Eq. (8) gives

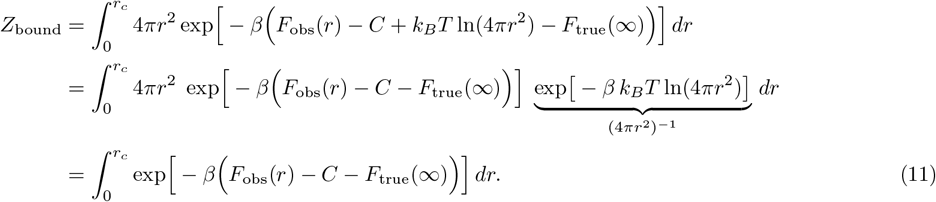

To eliminate the non–observable constants, we can evaluate Eq. (10) at the unbound plateau *r* = *r*_*c*_, where *F*_true_(*r*_*c*_) = *F*_true_(∞). This yields

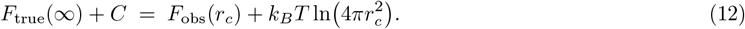

Inserting Eq. (12) into Eq. (11) gives

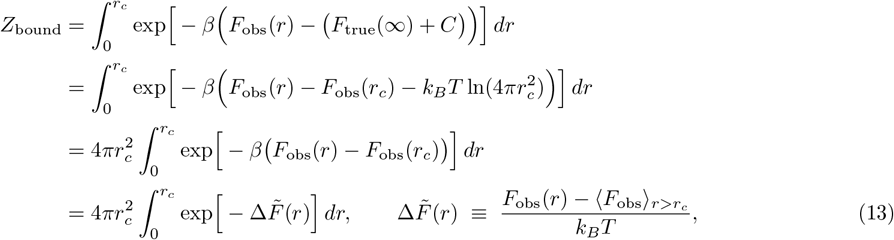

where we replace *F*_obs_(*r*_*c*_) by the mean value 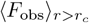 over the unbound plateau to reduce noise from fluctuations. Combining Eq. (13) with Δ*G* = −*k*_*B*_*T* ln(*ρ*°*Z*_bound_) reproduces Eqs. (2) and (3) in the main text.

##### Remarks.

(i) The Jacobian term 4*πr*^2^ (Eq. 8) cancels out exactly against the entropic contributions (Eq. 10), leaving the constant prefactor 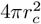 outside the integral. (ii) Any additive constant in *F*_obs_ is immaterial; only differences relative to the unbound plateau matter. (iii) *ρ*°*Z*_bound_ is dimensionless: *dr* has units of length, 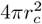 of length^2^. *Z*_bound_ has thus units of length^3^, while *ρ*° (1 M) has units of length^−3^, converting the “bound volume” (*Z*_bound_) to the standard state. We work in nanometers, hence *ρ*° = 0.602214 nm^−3^.

#### d. Thermal fluctuations upon binding

We also produce a reweighted histogram of the Root Mean Square Fluctuations (RMSF) of the targeted epitopes along the CV. In other words, we first bin all trajectory snapshots into 10 equal-width bins along the distance CV. Then, we align all snapshots within each bin using the Kabsch algorithm on the structured (non-IDR) residues, and compute the RMSF over the IDR residues only. Each snapshot *n* is assigned an importance weight

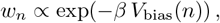

where the weights are normalized such that ∑ _*n*_ *w*_*n*_ = 1. The per-residue RMSF is

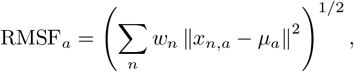

where *x*_*n,a*_ are the *C*_*α*_ position of the IDR amino acid (residue) *a* in snapshot *n* and *µ*_*a*_ = ∑ _*n*_ *w*_*n*_*x*_*n,a*_, i.e., the weighted mean position. For each bin, we report the RMSF taken as the mean and standard deviation across all the residues in the IDR. For all systems, this shows that by introducing the binder, i.e., at a low value of the CV, we reduce the thermal fluctuations of the IDR, and thus impose partial structuring. For some systems (P53-1, P53-2 or SUMO-1A), we can also see a wider distribution (i.e., greater standard deviation) of RMSF across the residues, potentially suggesting that near the transition, i.e., near *r*_*c*_, some parts of the IDR might fluctuate more than other parts, those likely still remaining bound to the peptide. We do not investigate this further. Lastly, additional analysis not displayed here did not show any significant increase in the secondary structure content imposed due to binding at low CV distances, such as alpha-helices or beta-sheets in the epitope.

#### e. Detailed results for the numerical convergence of the FES calculations

For each system, in Figures 9 to 19 we provide detailed plots describing the statistical sampling achieved in our simulations, as well as the convergence of the binding affinity Δ*G*. When plotting the FES, we only show the region below the plateau on the left-hand side, i.e., we crop high (positive) energies at low distances. The uncertainty of the FES (shown in the shaded region) is computed at each bin of the histogram by taking the standard deviation of the free energy surfaces retrieved from block analysis^25^ with 20 blocks. We checked, not shown here, that the Δ*G* estimate is invariant to the number of blocks, from 2 to 20 blocks. We show the convergence of Δ*G* as a function of simulation time. To demonstrate that our umbrella sampling simulations can properly reconstruct the FES, we compute the overlap of neighboring windows as:

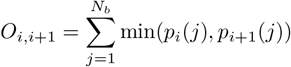

where *p*_*i*_(*j*) and *p*_*i*+1_(*j*) are the normalized histogram probabilities of window *i* and *i* + 1 in bin *j*, and *N*_*b*_ is the total number of histogram bins. We perform all binning at 100 bins, both here and during FES reconstruction in WHAM. Last, we also report the transition probabilities between replicas with a reasonable 20% acceptance rate between neighboring replicas.

**Figure 9.**
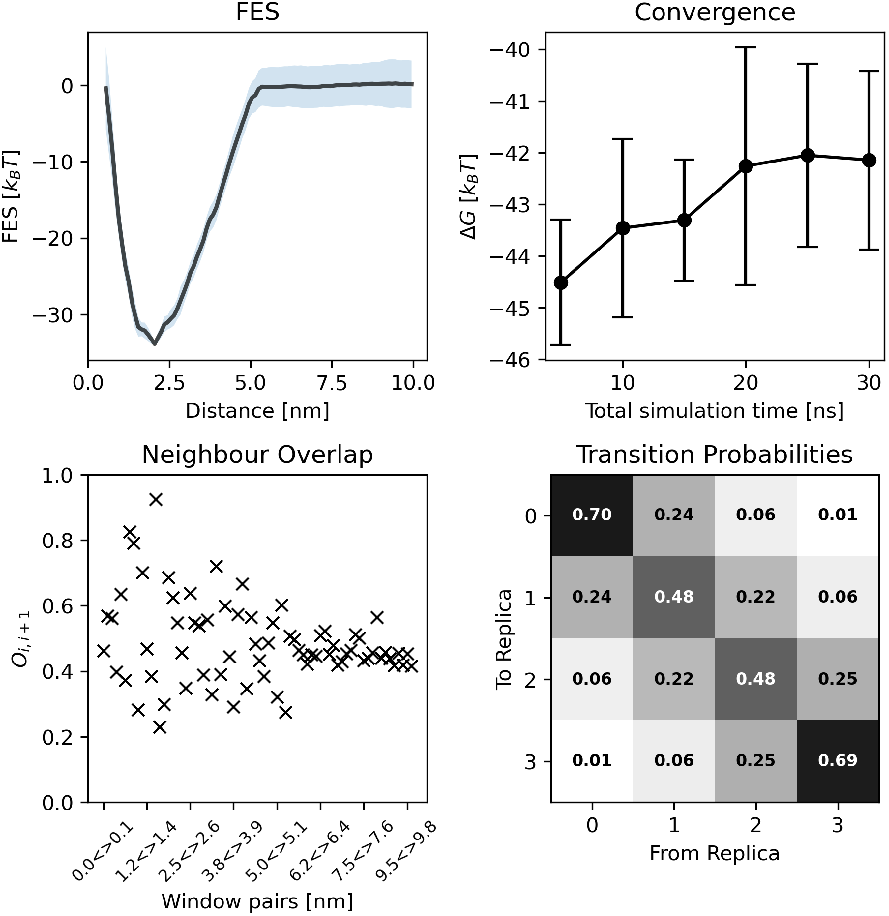
Free energy calculation details for ASYN-A.

**Figure 10.**
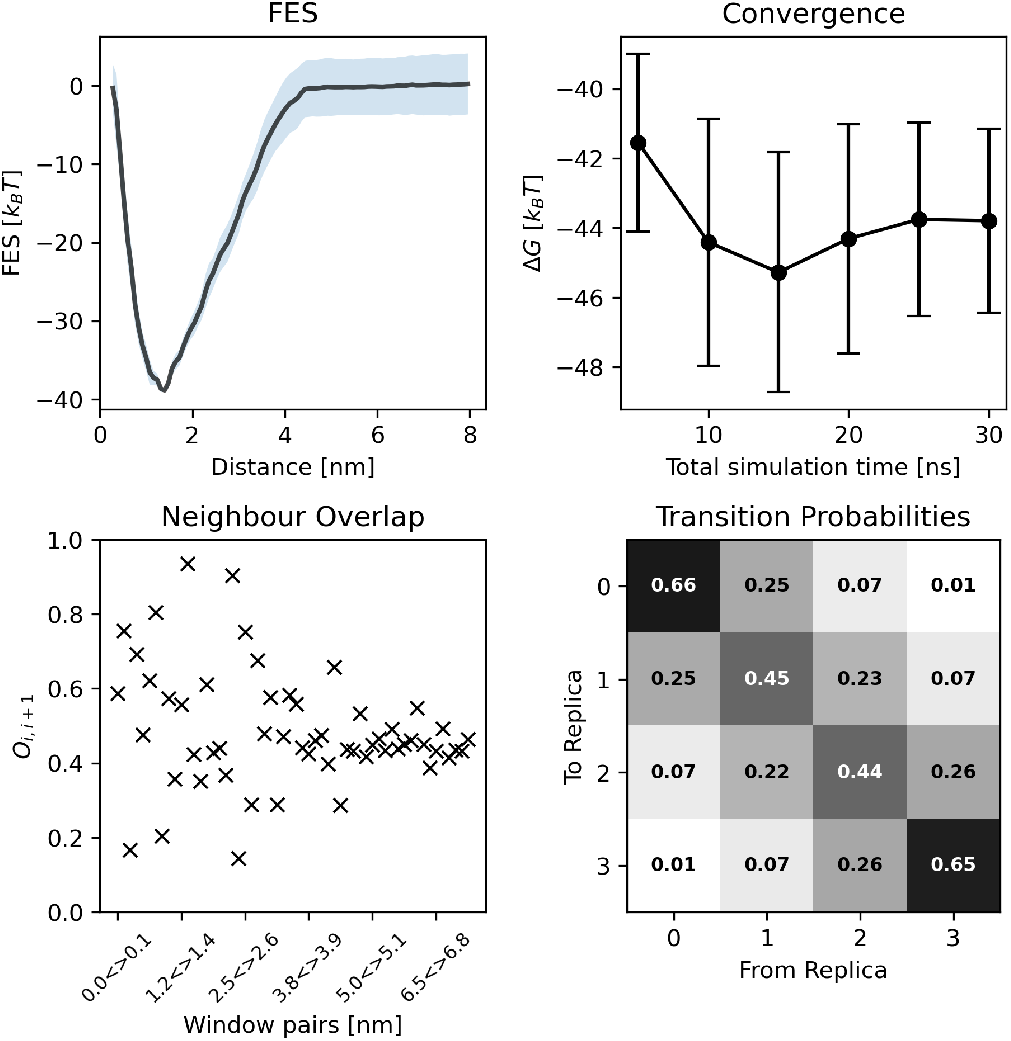
Free energy calculation details for ASYN-G.

**Figure 11.**
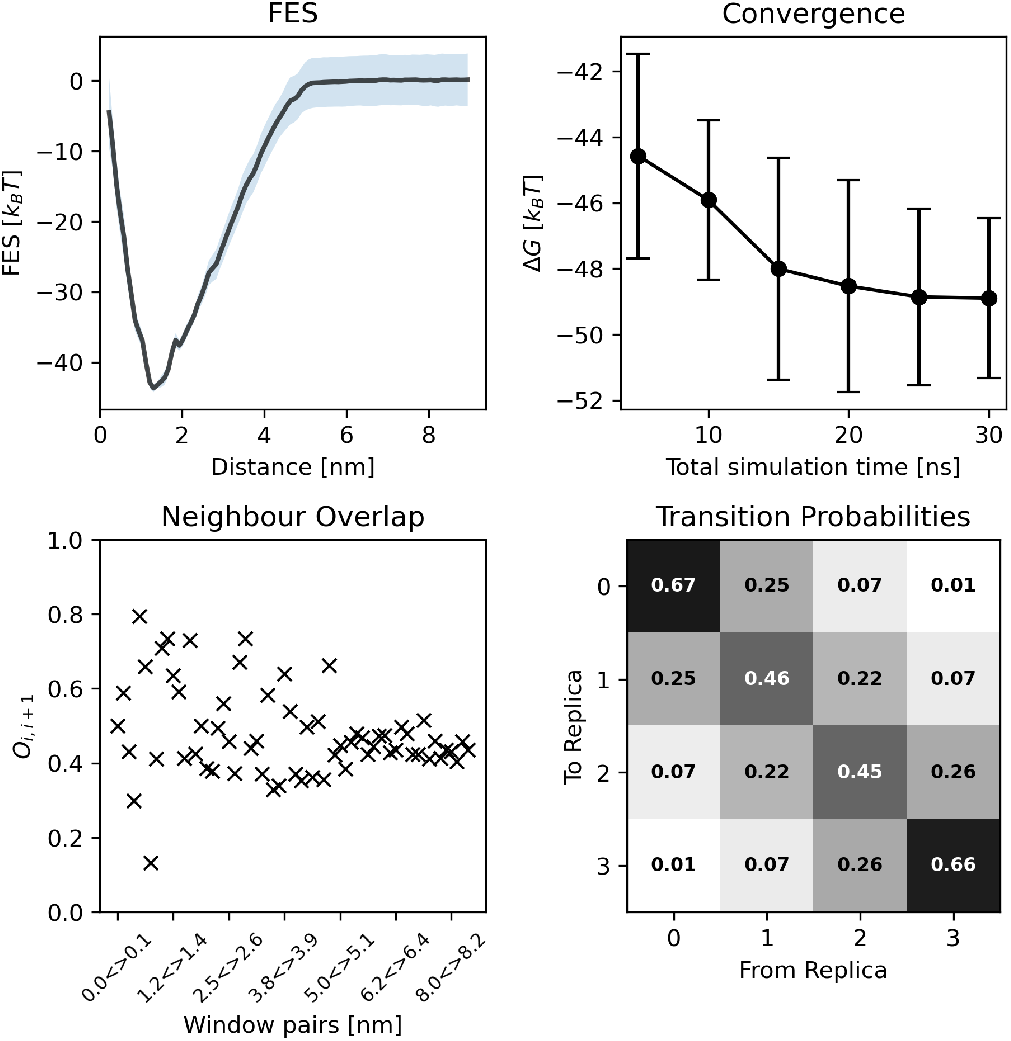
Free energy calculation details for CD28-A.

**Figure 12.**
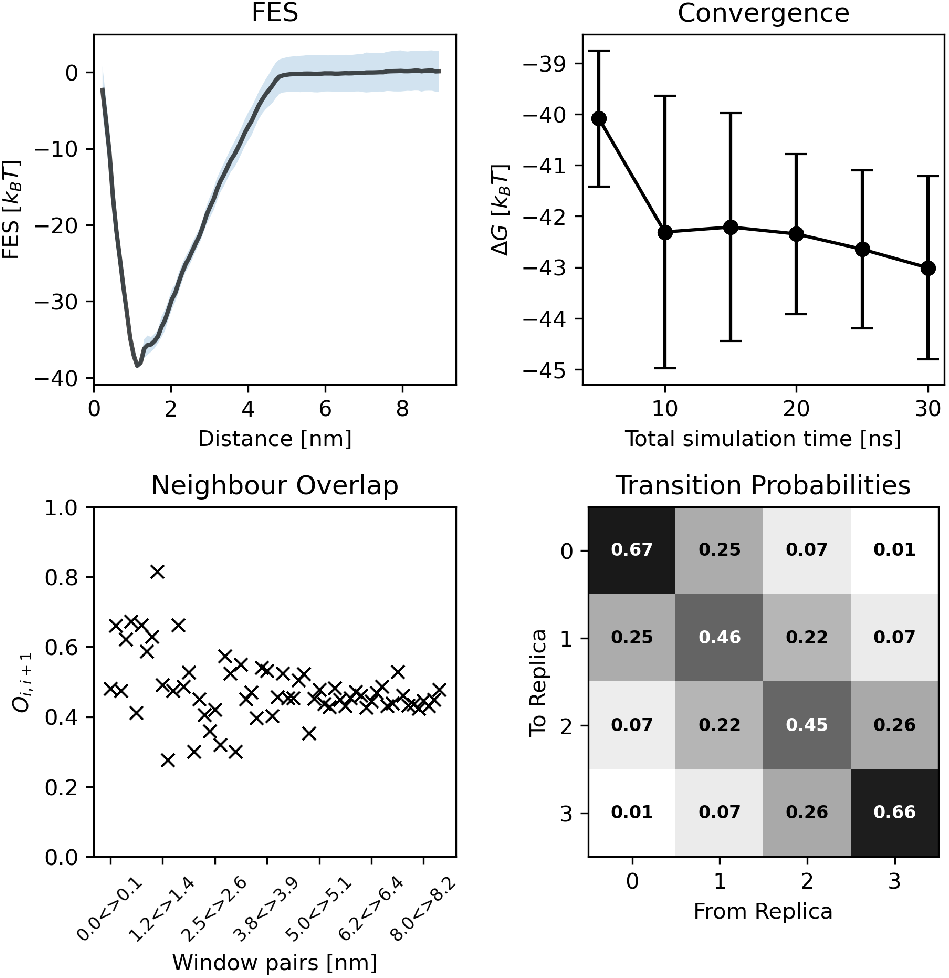
Free energy calculation details for CD28-B.

**Figure 13.**
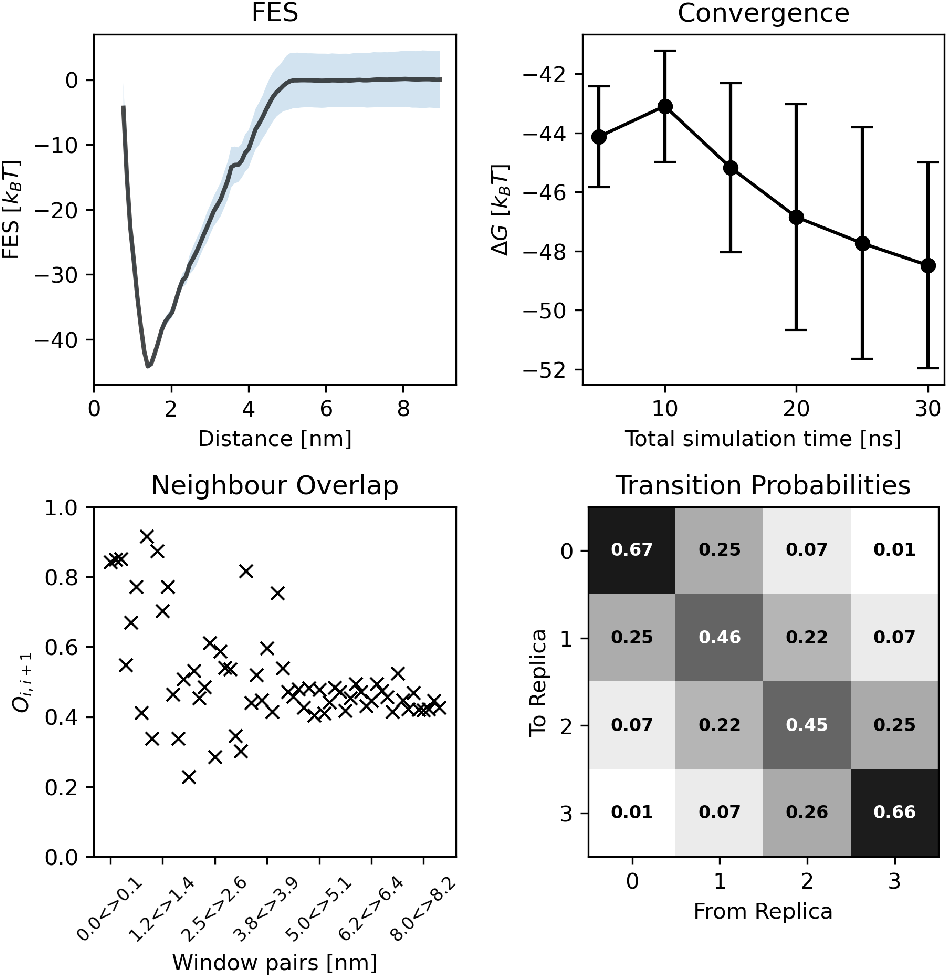
Free energy calculation details for CD28-G.

**Figure 14.**
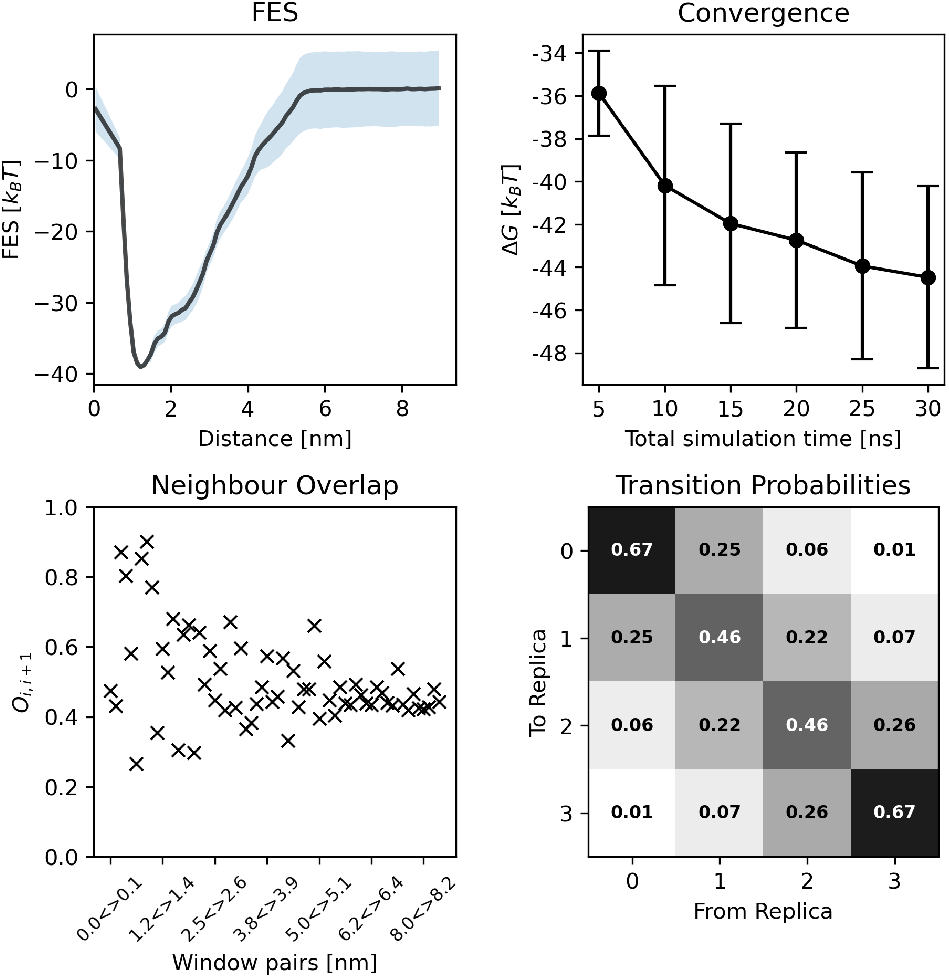
Free energy calculation details for CD28-P.

**Figure 15.**
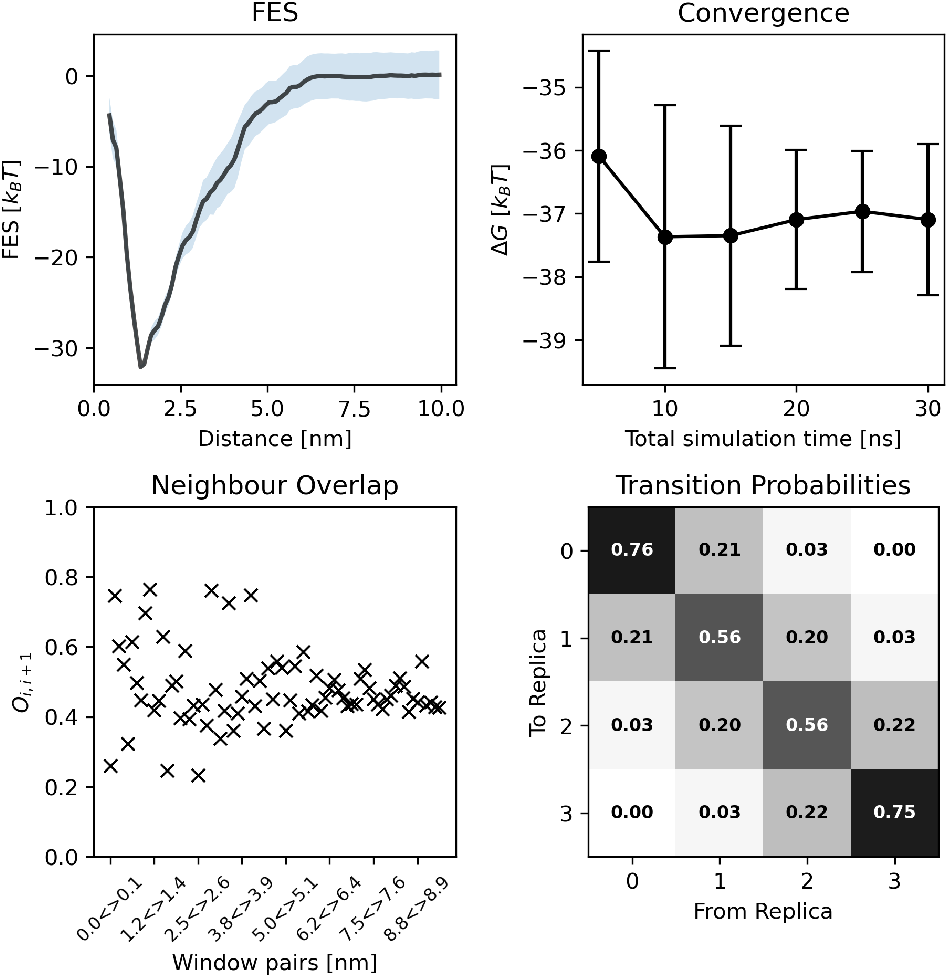
Free energy calculation details for P53-1.

**Figure 16.**
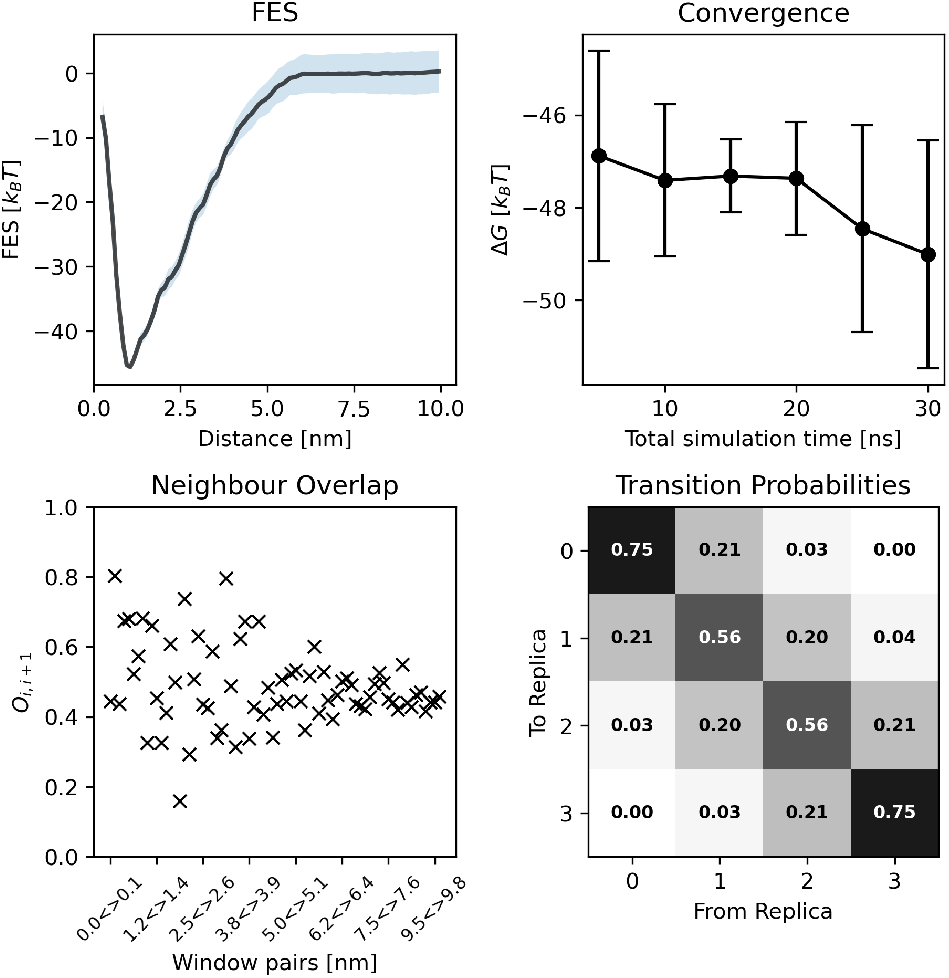
Free energy calculation details for P53-2.

**Figure 17.**
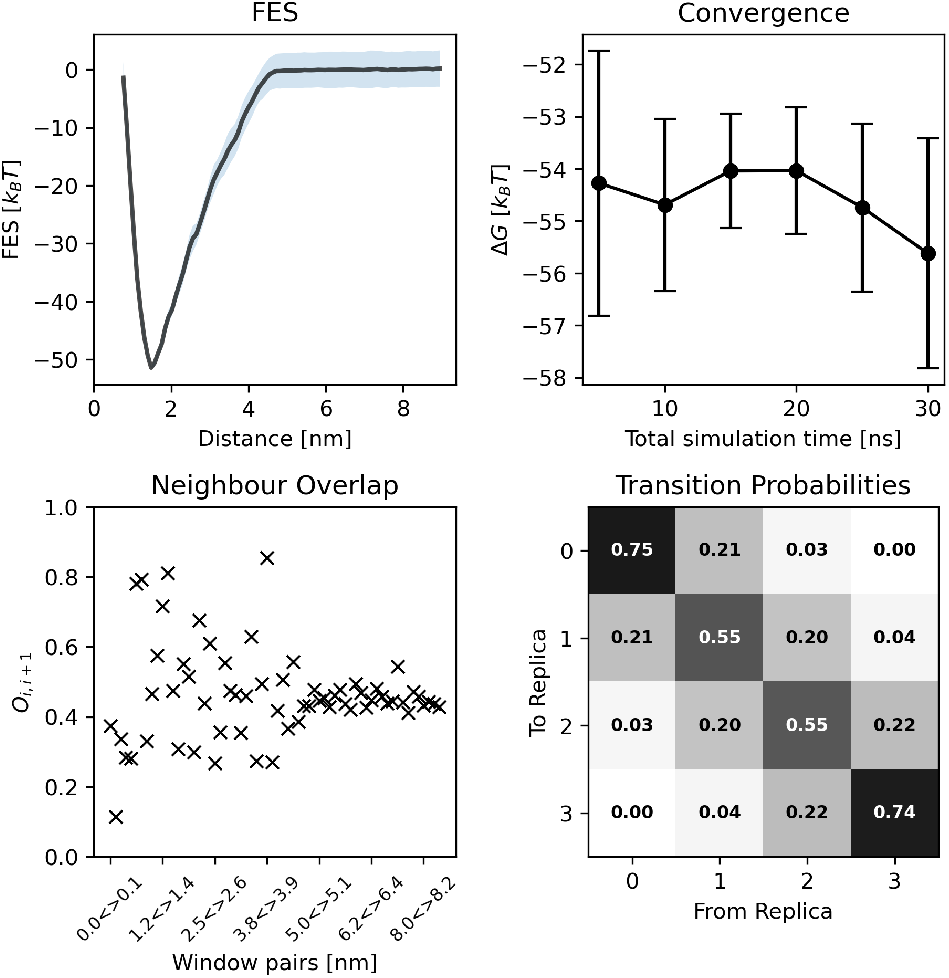
Free energy calculation details for P53-E.

**Figure 18.**
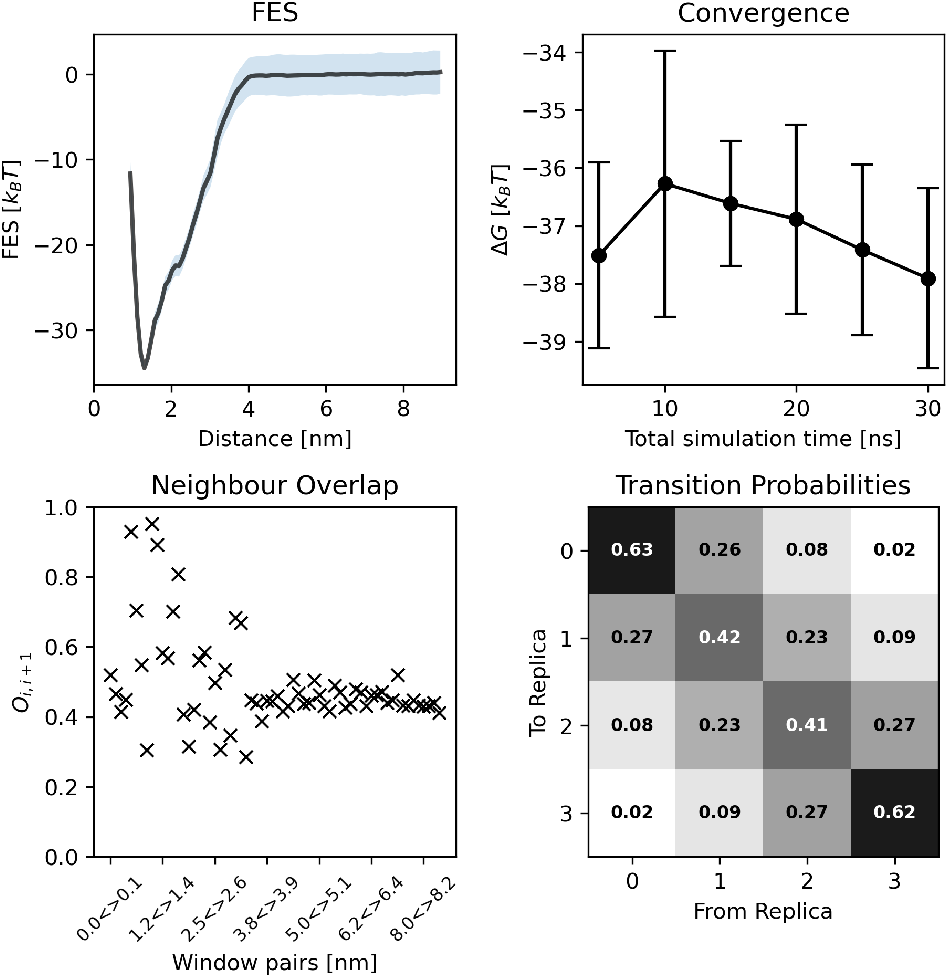
Free energy calculation details for SUMO-1A.

**Figure 19.**
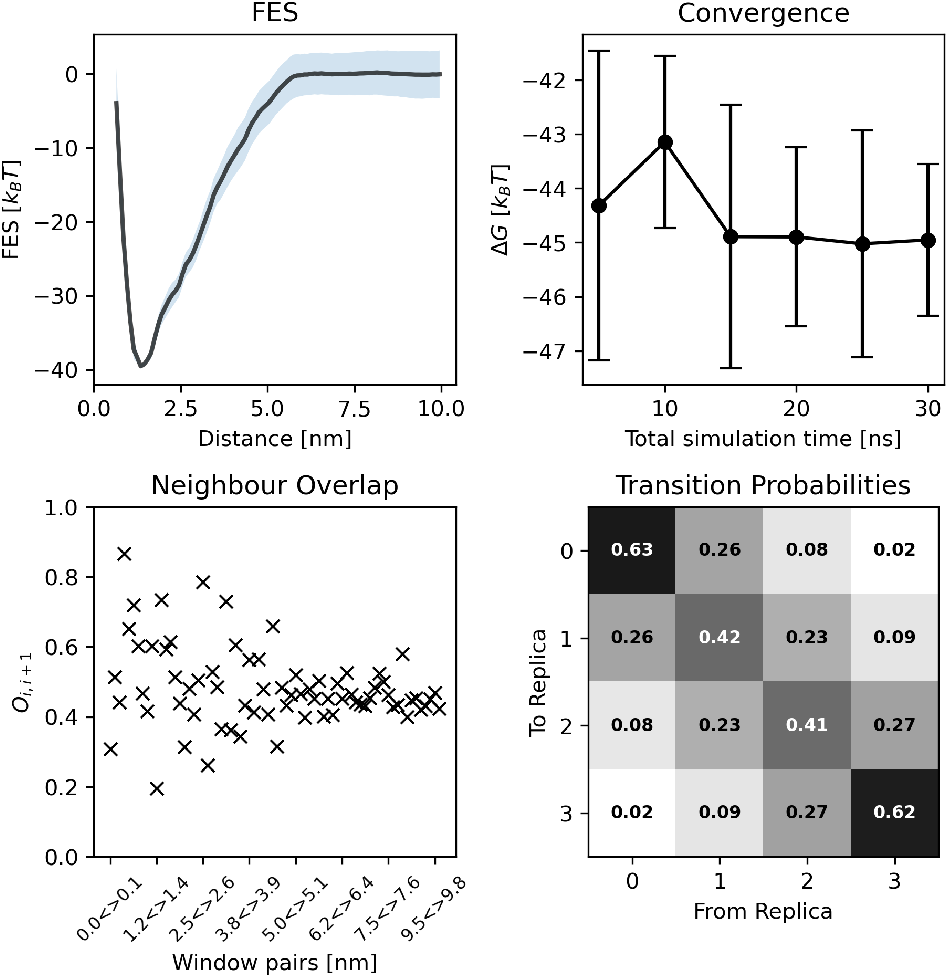
Free energy calculation details for SUMO-1C.

